# ProtCOVER: Family-Aware Conformal Prediction for Variant Effect Predictors on Unseen Protein Families

**DOI:** 10.64898/2026.11.13.766406

**Authors:** Wael Fahmy, Elena Krikun, Abedalrhman Alkhateeb

## Abstract

Protein language models and related variant effect predictors (VEPs) increasingly classify missense variants in proteins evolutionarily distant from those used for calibration. Split conformal prediction over variants is valid for its own target, a random variant dominated by well-studied families, but deployment needs coverage for a variant of a new family. On Prote-inGym clinical substitutions with five zero-shot predictors, Mondrian conformal comes within 0.005 of that target for four predictors, yet family-weighted pathogenic coverage is only 0.823– 0.873 at a 0.90 target for all five, without any shift. We present ProtCOVER, a conformal procedure that treats homology families, rather than individual variants, as the exchangeable unit. ProtCOVER (i) calibrates at the family level over label-free homology clusters, giving marginal coverage for a new family under family exchangeability; (ii) adapts to descriptor shift by calibrating conditionally on an estimated density ratio of label-free descriptors (sequence similarity, alignment depth and embedding distance; a variant reweights by the ratio instead), with finite-sample family-weighted coverage, valid as computed, whenever the true ratio is a nonnegative affine function of the estimate; and (iii) can abstain at low effective calibration support, which proved no better than random abstention. Under a shift of its descriptors, Prot-COVER’s mean worse-class shortfall below the target is 0.000, against 0.010 for group-weighted and 0.029 for hierarchical conformal prediction, with larger sets (about equal at matched coverage); on orphan families and held-out embedding regions it matches group weighting (0.002 against 0.003). In evaluations following pre-registered protocols, every shift-aware method met the target on new ClinVar variants of benchmark proteins, which ESM-1b separated better; on 12,225 proteins outside the benchmark, every method fell short for pathogenic variants with either of two predictors, ProtCOVER and a variant conditioned on the descriptors the least, within the 0.01 margin of group weighting.

## 1 Introduction

Zero-shot variant effect predictors (VEPs), from protein language models such as ESM-1b [38,9] to alignment-based models such as EVE and GEMME [16,23] and hybrids such as TranceptEVE and PoET [33,42], now rank missense variants with high accuracy on clinical benchmarks [32]. Clinical use, however, needs more than a ranking: variant interpretation under the guidelines of the American College of Medical Genetics and Genomics and the Association for Molecular Pathology (ACMG/AMP) [37] needs evidence whose error rate is known, and the ClinGen calibration of computational evidence [35] is genome-wide, while a single genome-wide calibration can hide large differences between genes [40].

Conformal prediction [44,1] offers distribution-free error control: a set over {benign, pathogenic} that contains the true label with probability at least 1 − *α*. Its guarantee rests on exchangeability between calibration and test data. For VEPs this assumption fails in two ways. First, variants are grouped into proteins and homologous protein families: a predictor’s accuracy differs from family to family, and families contribute very different numbers of labeled variants (from 2 to 741 per protein in our data). Pooling variants lets the largest families set the error rate, so the “90%” is a statement about variants from well-studied genes, not about a new protein. Second, the proteins a predictor is deployed on are often evolutionarily distant from, or have shallower alignments than, the proteins in the calibration set: a family-level distribution shift that is visible without labels. We address both with ProtCOVER, an uncertainty-quantification layer on top of a predictor’s scores that calibrates over homology families instead of variants: its output is a prediction set with controlled error, not a variant classification or an ACMG/AMP evidence strength.

We make three contributions.

– *Problem and evidence*. We formalize *family-weighted* coverage, the probability that a random variant of a new protein family is covered, and its class-conditional version for pathogenic variants, and show that variant-level calibration, which is valid for its own target, misses this one. On ProteinGym clinical substitutions, class-conditional (Mondrian) split conformal, the standard remedy for class imbalance, comes within 0.005 of its own variant-level target for four of five predictors yet covers only 0.823–0.873 of pathogenic variants per family for all five at a 0.90 target, with no distribution shift at all; pre-registered tests confirm this on ClinVar variants added after the benchmark (0.873 for ESM-1b) and on proteins outside it (0.778). Family-level calibration removes the failure when families are exchangeable.
– *Method and guarantees*. ProtCOVER calibrates at the family level and conditionally on a cross-fitted density ratio of label-free family descriptors; an optional gate can abstain when calibration support is low. Family-level pooling is not new: hierarchical conformal prediction (HCP) [25] is conformal risk control [2] with families as units. Nor is the form of our program: it is the conditional risk control of Blot et al. [7], which carries the conditional calibration of Gibbs et al. [18] to such losses. What Theorem 1 adds is a family-level construction that is valid in finite samples *as computed*: the test family’s labels are unknown, so its scores sit at +∞; the minimizer need not be unique, so the threshold is the largest optimal fitted value; the program can be unbounded, so the set is then {0, 1}; and each computed threshold is certified conservatively in exact arithmetic. The resulting family-weighted quantile regression has family-weighted coverage at least 1 − *α* under every family-level shift whose density ratio is a nonnegative function in the span of its features, and gives HCP’s threshold for constant features (up to ties). With the estimated ratio as a feature, ProtCOVER is valid under family exchangeability and when the estimated ratio is correct, and its loss under any other shift is bounded by the smaller of the bounds for HCP and for reweighting (Corollary 1).
– *Evaluation, including where the method does not help*. We test five predictors on five kinds of split (exchangeable; shifted descriptors; orphan families and held-out embedding regions, both defined without the descriptors; and a shift in predictor accuracy that violates our assumptions), with robustness checks of the base model, the descriptors and the family definition, and two external evaluations that follow pre-registered protocols, one on 12,225 proteins outside the benchmark with two predictors. Under descriptor shift ProtCOVER has the smallest worse-class shortfall below the target; on orphan families and held-out regions it matches group weighting, and it never exceeds a baseline’s shortfall by more than 0.01, at the price of larger sets under shift. When the predictor’s accuracy shifts, in a stress test and on the new proteins, every method under-covers, ProtCOVER’s conditional versions least but within the margin of group weighting; the temporal evaluation does not separate the shift-aware methods; and the optional gate is a negative result: its support-based abstention does not beat random abstention.

## 2 Related Work

### Conformal prediction

Split conformal prediction [44,26] gives marginal coverage under exchangeability, and Mondrian versions calibrate within categories [43,12]. Under covariate shift, weighted conformal prediction uses a known or estimated likelihood ratio [41,27], related methods handle label shift, non-exchangeability, proximity and multiple sources [36,4,20,10,29], and group-weighted conformal prediction (GWCP) [5], our main baseline, handles shifts in the proportions of known groups. Conditional calibration gives coverage over a class of covariate shifts by quantile regression with the test point imputed [18], related methods give group-conditional coverage [22,3], and conformal risk control and its conditional version extend these to bounded monotone losses [2,7], all for exchangeable points. For grouped data, Dunn et al. [14] proposed pooling and subsampling for a new group, Lee et al. [25] proved finite-sample guarantees for pooling (HCP), and later work uses a few observations from the test group [30]; Duchi et al. [13] calibrate one coverage threshold per environment, and macro-coverage [6] averages coverage over classes, in the spirit of our family weighting. In protein science, conformal methods have handled the feedback covariate shift of sequence design [15] and mining protein databases for function [8]. Table 1 compares the guarantees.

**Table 1.**
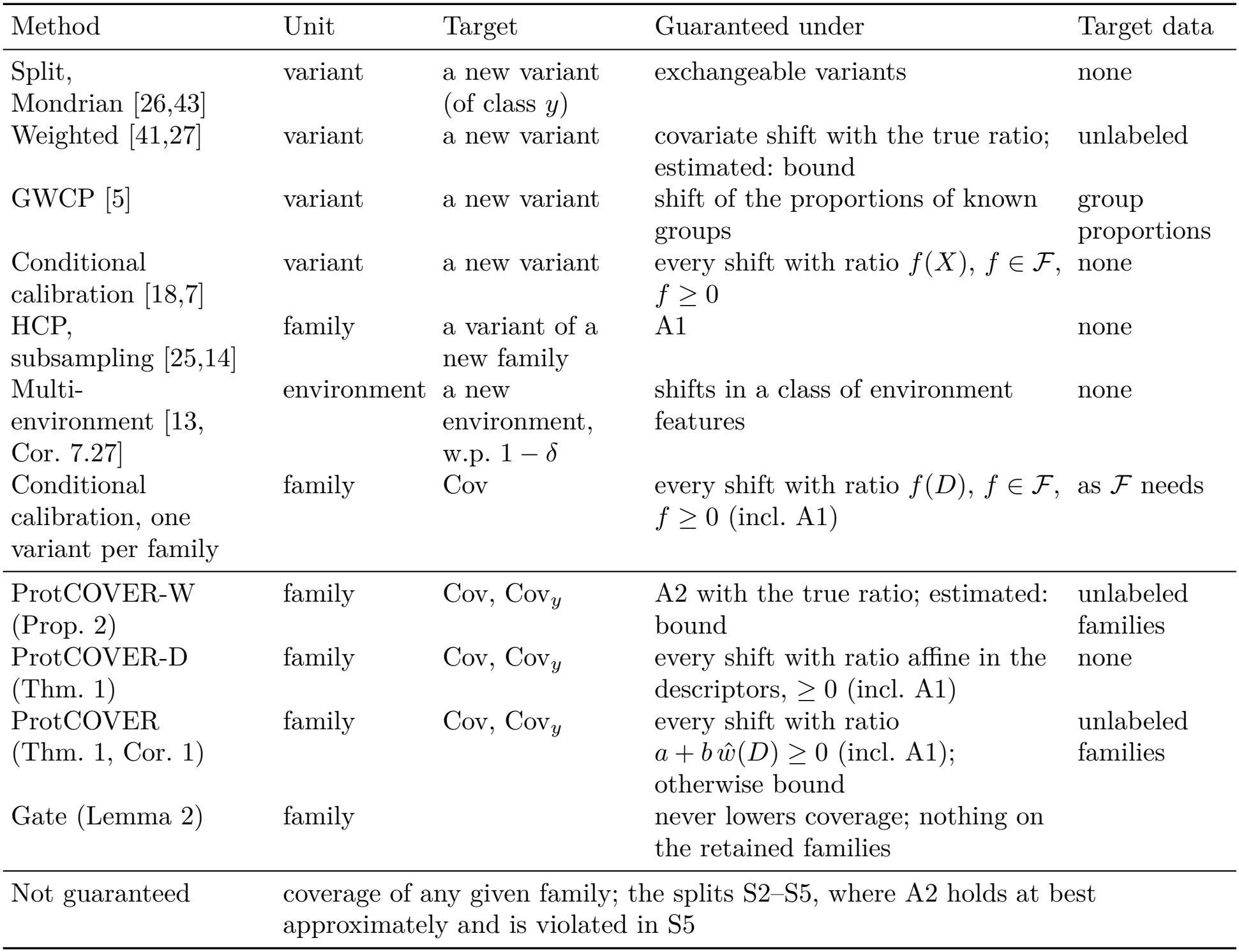
Guarantees of related methods and of ProtCOVER’s versions: finite-sample lower bounds of 1 − *α* on the target under the stated conditions, unless noted (repeated subsampling: 1 − 2*α*; multi-environment: with probability 1 − *δ*); “bound”: reduced by a total-variation distance that unlabeled data cannot estimate. ProtCOVER’s versions hold for Cov and Cov_*y*_ (Proposition 3) and, with cross-fitted ratios, conditionally on each fold’s ratio (Lemma 1).

| Method | Unit | Target | Guaranteed under | Target data |
|---|---|---|---|---|
| Split, Mondrian [26,43] | variant | a new variant (of class *y*) | exchangeable variants | none |
| Weighted [41,27] | variant | a new variant | covariate shift with the true ratio; estimated: bound | unlabeled |
| GWCP [5] | variant | a new variant | shift of the proportions of known groups | group proportions |
| Conditional calibration [18,7] | variant | a new variant | every shift with ratio *f*(*X*), *f* ∈ ℱ, *f* ≥ 0 | none |
| HCP, subsampling [25,14] | family | a variant of a new family | A1 | none |
| Multi-environment [13, Cor. 7.27] | environment | a new environment, w.p. 1 − *δ* | shifts in a class of environment features | none |
| Conditional calibration, one variant per family | family | Cov | every shift with ratio *f*(*D*), *f* ∈ ℱ, *f* ≥ 0 (incl. A1) | as ℱ needs |
| ProtCOVER-W (Prop. 2) | family | Cov, Cov<sub>*y*</sub> | A2 with the true ratio; estimated: bound | unlabeled families |
| ProtCOVER-D (Thm. 1) | family | Cov, Cov<sub>*y*</sub> | every shift with ratio affine in the descriptors, ≥ 0 (incl. A1) | none |
| ProtCOVER (Thm. 1, Cor. 1) | family | Cov, Cov<sub>*y*</sub> | every shift with ratio *a* + *b* *ŵ*(*D*) ≥ 0 (incl. A1); otherwise bound | unlabeled families |
| Gate (Lemma 2) | family |  | never lowers coverage; nothing on the retained families |  |
| Not guaranteed | coverage of any given family; the splits S2–S5, where A2 holds at best approximately and is violated in S5 |  |  |  |

### Calibration of variant effect predictors

ClinGen calibrated computational scores to evidence strengths for the codes PP3 (supporting pathogenicity) and BP4 (supporting benignity) [35]; later work found that genome-wide calibration masks heterogeneous performance across genes [40] and proposed gene- and domain-aware calibration [11], and RaCoon calibrates ESM-1b at the residue level [34]. These improve point probabilities within subgroups seen in training, without distribution-free guarantees, and condition on gene, domain or residue attributes rather than on homology support relative to the calibration data. Kermut [19] provides calibrated uncertainty for supervised variant effect regression on deep mutational scans. Selective classification [17] motivates our risk-coverage evaluation of abstention.

## 3 Problem Setup

### Families and scores

A protein *family* is a cluster of homologous wild-type sequences obtained without labels (Section 6.1). Family *k* is the random object 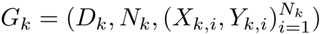, where *X*_*k,i*_ describes a missense variant, *Y*_*k,i*_ ∈ {0, 1} is its label (1 = pathogenic), and *D*_*k*_ ∈ *D* is a *label-free descriptor* of the family: any function of its sequences, alignments, predictor scores, and of the fixed set of training families, but not of labels. A predictor score is turned into 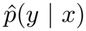 by a base model fitted on separate training families, which we treat as fixed. The nonconformity score is 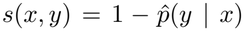 and *S*_*k,i*_ = *s*(*X*_*k,i*_, *Y*_*k,i*_). We observe calibration families *G*_1_, …, *G*_*K*_ with labels and a test family *G*_*K*+1_ whose descriptor *D*_*K*+1_ and variants *X*_*K*+1,*i*_ are observed but whose labels are not.

### Coverage targets

The unit of sampling is the family: a new family is drawn from the family-level law (of *G*_*K*+1_), and the question is how often its variants are covered. For a set-valued predictor *C* : *X* → 2^{0,1}^, the per-family coverage of the test family is 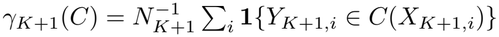, and our primary target is *family-weighted coverage*

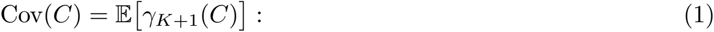

draw a new family, then one of its variants uniformly at random; Cov is the probability that this variant is covered. Every family counts once, whatever its number of variants. For example, among ten test families, one of 500 variants and nine of 5, the large one carries over 90% of the weight in variant-weighted coverage, which pools the variants (reported, not targeted), but a tenth in Cov. Its class-conditional version for label *y* is 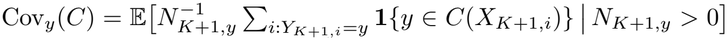, where *N*_*k,y*_ counts label-*y* variants of family *k*; it averages over the families that have label-*y* variants, not over all families. For a predictor used to flag pathogenic variants, Cov_1_ (pathogenic coverage) is the quantity of interest: 1 − Cov_1_ is the family-weighted *pathogenic miss rate*, the rate at which a pathogenic variant’s set excludes the pathogenic label. Coverage of each family cannot be guaranteed without assumptions on how families differ [43].

### Assumptions

We use two alternative assumptions; each result states which one it needs. **A1** (*Family exchangeability*.) (*G*_1_, …, *G*_*K*+1_) is exchangeable.

**A2** (*Family-level covariate shift*.) 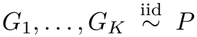 and, independently, the test family *G*_*K*+1_ and any further unlabeled target families are i.i.d. from *Q*, where *Q* ≪ *P* and 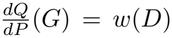 for a measurable *w* : *D* → [0, ∞). Equivalently, the descriptor’s law under *Q* is absolutely continuous with respect to its law under *P*, and the conditional law of a family’s variants and labels given its descriptor is the same under *P* and *Q*: only the distribution of descriptors shifts.

A1 holds when calibration and test families are drawn by a random split of a pool of families, regardless of dependence between families. It does not require exchangeability of variants within a family, because Cov averages over all variants of the test family (this is weaker than the hierarchical exchangeability of [25], which targets one variant of the new group). A2 formalizes deployment on families that differ from the calibration families in label-free ways, such as being evolutionarily distant or having shallow alignments; it fails when the predictor’s accuracy shifts in a way that the descriptors do not see. Our shifted splits (S2–S5, Section 6.2) draw test families without replacement from one finite pool, and S3–S5 select them by properties (orphan status, embedding region, predictor accuracy) that the descriptors do not determine. A2 therefore holds at best approximately in S2 and not in S3–S5, where the distances of Corollary 1 can be close to one, so those splits only measure robustness; the simulations of Appendix C check the case the theory covers.

## 4 ProtCOVER

ProtCOVER has four components: (i) family-level calibration; (ii) adaptation to descriptor shift; (iii) an optional support gate; and (iv) class-conditional versions of each. Its default (ii) is conditional calibration with features (1, *ŵ*), for an estimated density ratio *ŵ*. Two variants replace it: *ProtCOVER-W* reweights the families by *ŵ*, and *ProtCOVER-D* conditions on the descriptors instead of *ŵ*.

### (i) Family-level calibration

Let 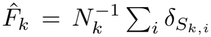 be the empirical distribution of calibration scores of family *k*. Given nonnegative family weights *ω*_1_, …, *ω*_*K*+1_, set 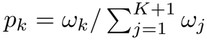 and

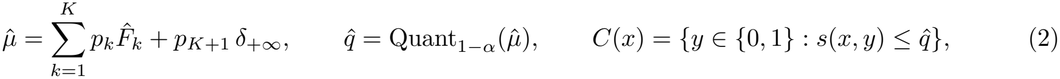

where Quant_*τ*_ (*µ*) = inf{*u* : *µ*((−∞, *u*]) ≥ *τ*}. With *ω* ≡ 1 every family, calibration or test, has the same total weight, spread evenly over its variants. The test family’s weight sits at +∞ because its labels, and hence its scores, are unknown: the guarantee holds for the threshold computed with its true scores, and moving them to +∞ can only raise the threshold. This is the hierarchical conformal prediction (HCP) threshold of [25], which equals the CDF pooling of [14] up to the level adjustment.

### (ii) Adapting to descriptor shift: conditional calibration on an estimated density ratio

Equation (2) uses one threshold for every test family. ProtCOVER lets the threshold depend on the test family’s label-free descriptor through a feature map *Φ* : *D* → ℝ^*p*^ with *Φ*_1_ ≡ 1, spanning ℱ = {*Φ*(·)^⊤^*b* : *b* ∈ ℝ^*p*^}. With *τ* = 1 − *α* and the pinball loss *ρ*_*τ*_ (*u*) = max{*τu*, (*τ* − 1)*u*}, let

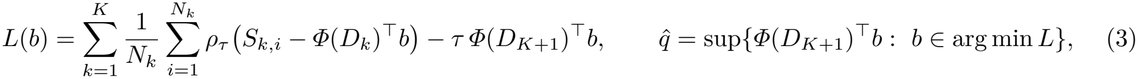

with 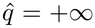 if *L* is unbounded below or if the fitted value *Φ*(*D*_*K*+1_)^⊤^*b* is unbounded over arg min *L*, and 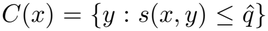. This is a family-weighted quantile regression in which, as in (2), each calibration family has total weight one and the test family’s scores sit at +∞ (the last Lterm); with *Φ* ≡ 1 it is the HCP threshold up to ties (Appendix A). *L* is bounded below iff −*τ Φ*(*D*_*K*+1_) = ∑ _*j*_ *a*_*j*_*Φ*(*D*_*k*(*j*)_) for some weights (*τ* − 1)*/N*_*k*(*j*)_ ≤ *a*_*j*_ ≤ *τ/N*_*k*(*j*)_ on the calibration scores *j*, of families *k*(*j*); otherwise the query’s features cannot be balanced by the calibration families, for example when its estimated ratio far exceeds theirs, and the set is {0, 1}: a built-in abstention. The default feature map is *Φ* = (1, *ŵ*), where *ŵ* is an estimate of the density ratio *w* = *dQ/dP* of descriptors, so the threshold can rise for families that are rare among the calibration families and frequent among the target families. Of the two variants, *ProtCOVER-W* instead reweights the families in (2) with *ω*_*k*_ = *ŵ*(*D*_*k*_), *k* ≤ *K* + 1 (computable because *D*_*K*+1_ is label-free), and *ProtCOVER-D* uses the standardized descriptors other than length as features and needs no target families.

The ratio comes from a logistic classifier separating calibration from target families on standardized descriptors, 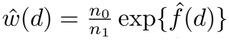 with 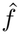 the fitted log-odds and *n*_0_, *n*_1_ the class sizes, rescaled to mean one over the fitting calibration families and clipped to [1*/*20, 20]. The descriptors measure sequence similarity (maximum identity to any training protein), alignment depth (log(1 + *N*_eff_) and log of *N*_eff_ per position, *N*_eff_ the effective number of sequences in the alignment) and embedding distance (to the nearest training protein, from ESM2 [28]), with log length (Appendix B). *Cross-fitting:* a third of the calibration families fits *ŵ* and the rest calibrate; the target families form five folds, and the ratio for queries in fold *j* is fitted against the target families outside fold *j*, so each fold’s guarantee is conditional on its own ratio (Lemma 1).

#### Computation

We solve (3) through its dual linear program, which has one bounded variable per calibration score but only *p* equality constraints; test families change only the right-hand side, so HiGHS [21] re-solves each from the previous basis (a few milliseconds). To obtain a value at least the supremum in (3) when arg min *L* is not a single point, the test term’s weight is raised from *τ* to *τ* +10^−4^: every exact minimizer of the perturbed program has a fitted value at least the supremum, and an unbounded perturbed program gives +∞ (Appendix A). Solvers are optimal only within tolerances, so the returned basis is certified optimal for the perturbed program in exact rational arithmetic (a cold re-solve, then an exact dual simplex, as fallbacks), and the fitted value is rounded up. The computed sets therefore contain those of (3) and keep every guarantee by Lemma 2 (Appendix B reports the checks).

### (iii) Support gate

The gate returns {0, 1} for every variant of a query family (it abstains) when few calibration families resemble it, measured by a kernel effective sample size (ESS) of the support descriptors (maximum identity, alignment depth, embedding distance) below *m*, or when the ratio puts more than *π*_max_ of ProtCOVER-W’s quantile mass on the query itself; its parameters are tuned on an inner split of the training families (Appendix B). It only enlarges sets, so results for ProtCOVER and its variants are without it.

### (iv) Class-conditional versions

Every component has a Mondrian version: for each label *y*, the threshold 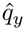 is computed from label-*y* calibration variants only (weights 1*/N*_*k,y*_; families with *N*_*k,y*_ = 0 dropped), and 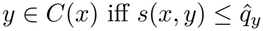. These target Cov_0_ and Cov_1_ separately and are the versions we compare.

## 5 Guarantees

All results are finite-sample and hold for any fixed score function, so they do not depend on the quality of the base model or of the predictor; proofs are in Appendix A. Each concerns the expected coverage of a random variant of a new family, not the coverage of every family. Table 1 lists what each covers.

### Proposition 1

**(Family-level calibration)**

*Under A1, the set* (2) *with ω* ≡ 1 *satisfies* Cov(*C*) ≥ 1 − *α*. Proposition 1 is Theorem 1 of [25] with the test point replaced by a uniformly random variant of the test family, which removes the need for exchangeability within families.

### Theorem 1

**(Coverage over a class of family-level shifts)**

*Let P be a law of families and Φ fixed. For f* ∈ ℱ *with f* (*D*) ≥ 0 *P -almost surely and* 0 *<* E_*P*_ *f* (*D*) *<* ∞, *let Q*_*f*_ *be the family law with dQ*_*f*_ */dP* = *f* (*D*)*/*E_*P*_ *f* (*D*). *If G*_1_, …, *G*_*K*_ *are i*.*i*.*d. P and, independently, G*_*K*+1_ ∼ *Q*_*f*_, *the set* (3) *satisfies* Cov(*C*) ≥ 1 − *α. The same holds under A1 (f* ≡ 1; *the families need only be exchangeable). For fitted features such as* (1, *ŵ*), *the guarantee is conditional on the fit and relies on cross-fitting’s independence (Lemma 1)*.

### Corollary 1

**(Any family-level shift)**

*With G*_1_, …, *G*_*K*_ *i*.*i*.*d. P as in Theorem 1 and a test family with any law Q*, Cov(*C*) ≥ 1 − *α* − inf_*f*_ *d*_TV_(*Q, Q*_*f*_), *the infimum over the f of Theorem 1. If Φ* = (1, *ŵ*) *for a ratio estimate ŵ* ≥ 0 *independent of the families, with* 0 *<* E_*P*_ *ŵ <* ∞ *and* 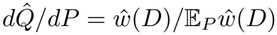, *then*

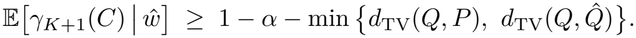

Here *d*_TV_(*Q, P*) bounds the loss of family-level calibration alone (the same argument applied to Proposition 1) and *d*_TV_(*Q*, 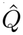) that of reweighting by *ŵ* (Proposition 2), so conditioning on the estimated ratio is never worse, in these bounds, than either. Both distances are unobservable, so the corollary motivates conditioning on both but certifies nothing under an arbitrary shift. Conditioning needs no total-variation term under family exchangeability, when *ŵ* is the true ratio up to scale, and for every shift whose ratio is a nonnegative function in the span of 1 and *ŵ*; with descriptor features (ProtCOVER-D), for every shift whose ratio is nonnegative and affine in the descriptors, without target families.

### What is new

Table 1 places these results among related guarantees. A family’s miss rate 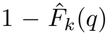 is a bounded loss that does not increase with the threshold, and family-weighted coverage is one minus its risk, which is why pooling families is the risk-control view applied to grouped data (Section 1); (3) applies conditional risk control to families with the test family’s loss imputed as one, and constant features give HCP’s threshold up to ties (Appendix A). For validity as computed, the largest optimal fitted value dominates the threshold of the program with the test family’s true scores (step (d) of the proof). This adapts Gibbs et al.’s Proposition 6, which imputes an upper bound for the unknown scores and assumes min-norm solutions; Blot et al. treat neither ties nor unbounded programs, and Section 4 computes the threshold exactly. Conditioning on the span of a constant and a cross-fitted ratio (Gibbs et al. note a single estimated shift as a one-element class) then gives Corollary 1. Applying Gibbs et al. to one random variant per family, the analogue of subsampling once [14], also gives family-weighted coverage over ℱ, because the subsampled pairs are exchangeable points, but discards all but one calibration variant of each family; we use it as a baseline. Duchi et al. [13, Cor. 7.27] regress one coverage threshold per environment on environment features, which gives a new family the conservative level (1 − *α*)(1 − *δ*), without pooling its variants.

#### Proposition 2

**(Reweighting variant)**

*Under A2, the set* (2) *with ω*_*k*_ = *w*(*D*_*k*_), *k* ≤ *K* + 1, *satisfies* Cov(*C*) ≥ 1 − *α. With ω*_*k*_ = *ŵ*(*D*_*k*_) *for ŵ* ≥ 0 *independent of the families with* 0 *<* E_*P*_ *ŵ*(*D*) *<* ∞ *and* 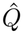 *as in Corollary 1*, E[*γ*_*K*+1_(*C*) | *ŵ*] ≥ 1 − *α* − *d*_TV_(*Q*, 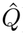).

Proposition 2 lifts weighted exchangeability [41] and its estimated-weight bound [27, Thm. 3] from variants to families.

#### Lemma 1

**(Cross-fitting)**

*For each fold j, the ratio ŵ*_*j*_ *is a function only of the fitting calibration families and of the target families outside fold j. With independent families (as under A2) it is therefore independent of the calibration families used in* (2) *and* (3) *and of every family in fold j, and Corollary 1 and Proposition 2 hold for the queries of fold j conditionally on ŵ*_*j*_. *If instead all non-training families are exchangeable, as under a random split, then conditionally on the families that fit ŵ*_*j*_ *the remaining calibration families and each family of fold j are exchangeable, and Theorem 1 with f* ≡ 1 *holds conditionally on ŵ*_*j*_.

#### Proposition 3

**(Class-conditional versions)**

*For y* ∈ {0, 1} *such that a family has N*_*y*_ *>* 0 *with positive probability under P and under the test law, the Mondrian versions of Propositions 1 and 2, Theorem 1, Corollary 1 and Lemma 1 hold with* Cov_*y*_ *in place of* Cov *and with Q, P*, *Q*_*f*_, 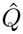 *conditioned on N*_*K*+1,*y*_ *>* 0.

#### Lemma 2

**(Enlargement)**

*If C*^′^(*x*) ⊇ *C*(*x*) *for all x almost surely, where C*^′^ *may depend on all observed data, then* Cov(*C*^′^) ≥ Cov(*C*) *and* Cov_*y*_(*C*^′^) ≥ Cov_*y*_(*C*).

The gate (iii) only replaces a set by {0, 1}, so by Lemma 2 it preserves every guarantee above, whatever its parameters; it gives no guarantee for the families it retains.

## 6 Experiments

### 6.1 Data, predictors, and families

We use the ProteinGym v1.3 clinical substitution benchmark [32]: 62,727 ClinVar [24] missense variants in 2,525 human proteins, 51.0% pathogenic, with a median of 12 labeled variants per protein (maximum 741). We take ProteinGym’s precomputed zero-shot scores for five unsupervised predictors: ESM-1b (protein language model), EVE and GEMME (alignment-based), TranceptEVE (hybrid), and PoET (retrieval-augmented language model), oriented so that higher means more pathogenic; our per-protein mean AUCs match ProteinGym’s published ones (0.892–0.920) to three decimals. Supervised clinical predictors, trained on ClinVar, are excluded.

Families are MMseqs2 [39] clusters of wild-type sequences at 30% identity and 80% bidirectional coverage: 2,081 families, 1,805 of them single proteins. Three alternative definitions serve as sensitivity checks (Appendix B): 50% identity (2,275 families), single-linkage components of significant homology (1,939), and, strictest, components of *any* significant hit (1,024, the largest with 867 proteins; a stress test for leakage, not a biological family). Descriptors are computed without labels (Section 4); a family is an *orphan* if no member has a significant hit at ≥ 30% identity outside the family (1,276 families).

### 6.2 Protocol

Each of ten seeds splits families into 40% training, 30% calibration and 30% test (832, 624 and 625 families). **S1** draws all three at random. **S2** keeps the S1 training families and samples test families with probability proportional to exp(−*βz*), *z* the mean of standardized maximum identity to training and log(1 + *N*_eff_), *β* ∈ {0.5, 1, 2}; it shifts the very descriptors that ProtCOVER and GWCP use, so it favors them by construction. Two shifts are defined without those descriptors, though they move them: **S3** draws test families from orphan families (mean maximum identity to training 0.048 against 0.236 for calibration families), and **S4** holds out whole regions of protein space, *k*-means clusters (*k* = 20) of the leading principal components of family-mean ESM2 embeddings, until the test set is full; a strict variant, **S4s**, holds out only whole regions (Appendix B). **S5**, a stress test that violates A2 on purpose, samples test families as S2 does, with weight exp(−*z*), *z* a family’s standardized within-family AUC averaged over predictors, so that predictors err more on them, a shift no label-free descriptor encodes. S4, S4s and S5 were declared before their first run.

The base model, fitted on training families only, is a logistic regression on the raw score, so that each variant is scored on its own, as in deployment; its test AUC is 0.872–0.916. A transductive variant adds the score standardized within its protein and the family’s number of scored variants as a descriptor; both are label-free but need the protein’s whole variant panel, and we report it as a robustness check. Baselines are variant-level split conformal (B1) and its class-conditional (Mondrian) version (B2); HCP pooling and Dunn et al.’s single and repeated subsampling (B3) [25,14]; GWCP (B4) [5] with nine descriptor bins and target proportions from the test families; and support-only and random abstention at the gate’s abstention rate (B5). ProtCOVER uses 416 of the calibration families in (2)–(3) and the rest to fit *ŵ*; ProtCOVER-D uses all calibration families, and the gate has a pre-declared 10% abstention budget. Test families are used only for evaluation, and only their label-free descriptors enter *ŵ*. Appendix B lists every setting.

#### Comparison criterion

Because family-level calibration can err in either class (for some predictors it over-covers pathogenic and under-covers benign variants), we compare class-conditional methods by the *worse-class shortfall* max{0, 1 − *α* − min(Cov_0_, Cov_1_)}, per predictor and setting, and treat differences below 0.01 as no difference, the criterion that we pre-registered for the external evaluation. The margin is statistical, not clinical: it lies between the median bootstrap half-width of a cell’s class-wise coverage (0.005) and the median seed-to-seed s.d. (0.013). Intervals are 95% family-bootstrap intervals that resample test families within each seed (paired across methods); they are conditional on the splits, and since all seeds resplit one pool of families, inference is limited to this benchmark. Unless stated otherwise *α* = 0.1.

#### External evaluations

Two evaluations calibrate on benchmark variants and test on ClinVar variants that played no part in development. Each follows a protocol pre-registered in the public repository before any method was run on its data, with amendments and post-hoc decisions explicitly documented (Appendix B). Both use the ESM-1b scores of Brandes et al. [9] and, as primary analysis, the raw-score base model (Appendix B). The *temporal* evaluation tests, in the benchmark’s splits, on 14,510 variants in 1,731 benchmark proteins that entered ClinVar after its snapshot: new variants and labels, not new proteins. The *new-protein* evaluation tests on 101,197 variants (9.4% pathogenic) in 12,225 human proteins outside the benchmark, clustered as for the benchmark into 8,579 test families (primary population) of which 4,448 have no detectable benchmark homolog (secondary); training families are each seed’s S1 draw and every other benchmark family calibrates. New proteins have no alignments, so the descriptors are alignment-free. A second predictor, ESM2 with 150M parameters scored by us on both sides [28,31], was pre-registered before any result.

### 6.3 Results

#### Variant-level calibration misses pathogenic variants per family

In S1 (Appendix Fig. 1), variant-level split conformal has variant-weighted coverage 0.883–0.899 and family-weighted coverage 0.871–0.903; the failure is in the pathogenic class. Mondrian split conformal (B2) attains variant-weighted pathogenic coverage 0.895–0.904 for four predictors yet covers only 0.823–0.873 of pathogenic variants per family for all five. The 5% of test families with the most pathogenic variants hold 50% of them, and their pathogenic variants are easier: under B2, families with fewer than five pathogenic variants have pathogenic coverage 0.781– 0.854 against 0.897–0.906 for families with at least twenty. For TranceptEVE even the variant-level target is missed (0.864; a family-level split does not make variants exchangeable), and its benign coverage per family is 0.802. Family-level class-conditional calibration attains 0.900–0.904 (pathogenic) and 0.897–0.902 (benign), as Proposition 3 guarantees in expectation, and ProtCOVER 0.902–0.904 and 0.898–0.908 with mean set sizes 1.171–1.405 (HCP 1.162–1.425): without shift, conditioning on the ratio costs little.

#### Under descriptor shift ProtCOVER has the smallest shortfall; on the other shifts it matches group weighting

We read S2, which is defined by the descriptors that ProtCOVER and GWCP use, separately from S3 and S4, which are not (Table 2). In S2, ProtCOVER has the smallest mean worse-class shortfall (0.000; GWCP 0.010, HCP 0.029), smaller than GWCP’s by more than the 0.01 margin in 4 of the 15 S2 cells (all at *β* = 2, where its sets are larger) and than HCP’s in 13. On S3 and S4 it is 0.002, against 0.003 for GWCP and 0.006 for HCP; paired intervals put ProtCOVER’s worse-class coverage above GWCP’s in 8 of 10 cells and below in 1 (differences −0.002 to +0.012), but the shortfalls are within the margin in every cell. Over all 25 shifted cells, ProtCOVER’s worst cell is 0.892 (HCP 0.820, GWCP 0.876), and its shortfall never exceeds a baseline’s by more than the margin. On S4, reweighting alone (ProtCOVER-W) falls short by 0.007 (GWCP 0.003), while ProtCOVER, which keeps the constant feature, meets the target in every S4 cell. The ordering holds at *α* = 0.05 and 0.2 (mean shortfall 0.000 and 0.002 for ProtCOVER, 0.005 and 0.011 for GWCP; Appendix Tables 12 and 13), at *α* = 0.05 with built-in abstention on up to 2.8% of test families.

**Table 2.** Worse-class shortfall below the 0.90 target, max{0, 0.9 − min(Cov_0_, Cov_1_)}, of class-conditional family-level methods at *α* = 0.1. Benchmark and temporal rows: mean (maximum) over the row’s predictor-setting cells, S2 pooling *β* ∈ {0.5, 1, 2}; a maximum of 0.000 means the target is met in every cell. New-protein rows: the single result of that predictor and population (the row on benchmark lengths is exploratory). S5 violates A2. Last two rows: mean set size over the benchmark’s shifted cells, at *α* = 0.1 and at pathogenic coverage 0.90 (interpolated).

|  |  |  | ProtCOVER |  |  |
|---|---|---|---|---|---|
| Evaluation | HCP | GWCP | -W | -D | **default** |
| *Benchmark, five predictors (development data)* |  |  |  |  |  |
| S2, descriptor shift | 0.029 (0.080) | 0.010 (0.024) | 0.001 (0.007) | 0.005 (0.016) | 0.000 (0.004) |
| S3, orphan families | 0.008 (0.017) | 0.003 (0.008) | 0.003 (0.007) | 0.001 (0.003) | 0.003 (0.008) |
| S4, held-out regions | 0.003 (0.013) | 0.003 (0.006) | 0.007 (0.013) | 0.004 (0.011) | 0.000 (0.000) |
| S4s, whole regions only | 0.005 (0.013) | 0.004 (0.009) | 0.002 (0.005) | 0.000 (0.001) | 0.000 (0.000) |
| S5, label-based shift | 0.122 (0.139) | 0.120 (0.138) | 0.118 (0.134) | 0.114 (0.130) | 0.116 (0.129) |
| *External ClinVar variants after the snapshot, ESM-1b, S2–S4 (pre-registered)* |  |  |  |  |  |
| Raw-score base model (primary) | 0.004 (0.020) | 0.000 (0.000) | 0.000 (0.000) | 0.000 (0.000) | 0.000 (0.000) |
| Transductive base model | 0.005 (0.023) | 0.000 (0.000) | 0.000 (0.000) | 0.000 (0.000) | 0.000 (0.000) |
| *Proteins outside the benchmark, alignment-free (pre-registered; all new families primary)* |  |  |  |  |  |
| ESM-1b, all new families | 0.036 | 0.034 | 0.032 | 0.025 | 0.027 |
| ESM-1b, no benchmark homolog | 0.083 | 0.079 | 0.077 | 0.048 | 0.062 |
| ESM-1b, benchmark lengths only | 0.021 | 0.019 | 0.017 | 0.011 | 0.012 |
| ESM2, all new families | 0.022 | 0.021 | 0.019 | 0.015 | 0.014 |
| ESM2, no benchmark homolog | 0.035 | 0.033 | 0.030 | 0.035 | 0.026 |
| *Mean set size, benchmark S2–S4* |  |  |  |  |  |
| At α = 0.1 | 1.228 | 1.241 | 1.262 | 1.252 | 1.294 |
| At matched Cov<sub>1</sub> = 0.90 | 1.266 | 1.259 | 1.263 | 1.253 | 1.268 |

#### The price is a more conservative operating point, not less efficient sets

ProtCOVER adapts how conservative it is to the shift: its mean worse-class coverage is 0.903 in S1 (HCP 0.900) but 0.906 over the shifted cells (GWCP 0.893, HCP 0.881; Appendix Fig. 2), so its sets under shift are larger; at matched pathogenic coverage the family-level methods need about the same set size (last two rows of Table 2; Appendix Fig. 3).

#### A label-based shift defeats every method, and the weights do not see it

In S5, every faLmily-levelLmethod falls to a worse-class coverage of 0.761–0.825 (Table 2), while the effective sample size 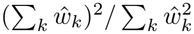 of the calibration weights is 0.98 of the number of families, as in S1 (0.99; S2 at *β* = 2: 0.26): the weights neither correct nor flag such a shift, and a weight diagnostic can show that a shift is visible, never that none exists.

#### Robustness, and method against information

The ranking under shift is unchanged with the transductive base model and, on S4s and under the three alternative family definitions, ProtCOVER’s mean shortfall is again at most GWCP’s and HCP’s (Table 2; Appendix Table 5). Without alignment depth, the descriptor that drives the S2 correction, the shift-aware methods fall further short, but ProtCOVER’s shortfall is still smaller than GWCP’s by more than the margin in 4 cells and larger in none. Conditional calibration on one random variant per family, which already attains family-weighted coverage, falls short by 0.002 over the shifted cells (ProtCOVER 0.001), with the same set size at matched coverage but coverage that varies more across seeds; richer feature classes are more conservative, and other ratio-fit shares change ProtCOVER’s shortfall by less than the margin (Appendix D). With ProtCOVER’s information, GWCP on ninths of *ŵ* falls short by 0.006 (its own bins: 0.007); with GWCP’s, conditional calibration on its bins falls short by 0.000 but abstains on up to 13.4% of a cell’s test families (ProtCOVER 0.3%; Appendix Table 6). The gain over GWCP comes mainly from conditioning, not the ratio’s information; conditioning on the ratio, not nine bins, keeps abstention rare.

#### New variants of benchmark proteins

The pre-registered hypotheses of the temporal evaluation were supported (Appendix B; H4’s rule was fixed after the run; Mondrian split covers 0.873 [0.864, 0.883] of pathogenic variants per family in S1), but ESM-1b separates these variants better (per-protein AUC 0.930, against 0.899 on their benchmark variants), and the data do not discriminate between the shift-aware methods: GWCP, ProtCOVER and its variants meet the target in every shifted setting; only HCP falls short (0.004).

#### On proteins outside the benchmark, every method under-covers pathogenic variants, the conditional versions the least

The pre-registered hypotheses were that Mondrian split covers less than 0.90 of pathogenic variants per family (U1), that ProtCOVER’s worse class covers at least 0.89 (U2), and that ProtCOVER’s shortfall exceeds neither HCP’s nor GWCP’s by 0.01 or more (U3) and is smaller than GWCP’s by more than 0.01 (U4). On the new proteins, Mondrian split under-covers pathogenic variants per family (0.778 [0.771, 0.786]; U1 supported; `docs/unseen_results.md`). Every family-level method falls short of the target in the pathogenic class (Table 2) while over-covering benign variants (0.965 to 0.967); ProtCOVER’s worse class covers 0.873, so U2 is not supported. ProtCOVER’s shortfall (0.027) exceeds neither HCP’s (0.036) nor GWCP’s (0.034) by the margin (U3 supported), but is not smaller than GWCP’s by more than it (U4 not supported); without detectable homologs the gap to GWCP exceeds it (0.062 against 0.079). ProtCOVER-D, outside the hypotheses, falls short least (0.025; without homologs 0.048). At matched coverage ProtCOVER’s sets are no larger (1.366, GWCP 1.372). The density ratio barely departs from one (effective sample size 0.96 of the calibration families), while ESM-1b separates the new proteins’ variants worse (per-protein AUC 0.873 against 0.892). The 223 families with a protein longer than any benchmark protein carry much of the shortfall (without them, exploratory, ProtCOVER’s is 0.012), and neither the ratio nor the gate flags them (ProtCOVER abstains on 0.0% of families). The ratio includes log length, but that does not guarantee detecting proteins outside the calibration range: benchmark proteins are longer on average, the fitted length coefficient is negative in every seed, and logistic extrapolation places these families near the bottom of *ŵ* (median rank 1.5% among new families, mean *ŵ* 0.56 against 1.10), none of them clipped. The gate’s descriptors do not include length: 14.1% of these families lie in the lowest decile of its ESS, and, tuned on the benchmark, it abstains on no new family. These are plausible explanations from exploratory diagnostics, not proven causes. With ESM2 (per-protein AUC 0.795 on the new proteins, 0.808 on the benchmark), the four verdicts are the same (Mondrian split 0.850) and the conditional versions again fall short least (ProtCOVER 0.014, ProtCOVER-D 0.015; HCP 0.022, GWCP 0.021), but without homologs ProtCOVER’s gap to GWCP is within the margin (0.026 against 0.033) and ProtCOVER-D falls short as much as HCP (0.035).

#### The support gate preserves coverage but does not improve selective risk

We keep the gate as an informative negative experiment, not a benefit of ProtCOVER. Abstaining first, on ProtCOVER-W’s sets, on the families with the lowest ESS lowered the area under the risk-coverage curve (AURC) relative to random abstention in the majority of seeds in 13 of 30 cells and raised it in 14 (Appendix Fig. 4); ordering by maximum identity alone did better (lower in 23 cells, higher in 6), with small mean differences from random (−0.011 to +0.017). Support descriptors locate the families whose coverage needs correcting, which conditional calibration already does; once coverage is controlled they predict little of where the predictor errs.

## 7 Limitations

### What the guarantees assume

Coverage of at least 1 − *α* needs exchangeable families (A1), or a family-level shift whose density ratio is a nonnegative function in the span of the features, with an estimated ratio fitted on families independent of those it calibrates (Lemma 1); the class-conditional versions hold for families that contain the class. Every guarantee is marginal over families: ProtCOVER certifies no particular protein, and the coverage of a small family is dominated by binomial noise. Under any other shift, Corollary 1 bounds the loss by total-variation distances that unlabeled data cannot estimate; when a family’s scores and labels given its descriptors differ between calibration and deployment (A2 fails, as in S5 and apparently on the new proteins), every method under-covers and the ratio gives no warning. Including log length does not guarantee flagging test families outside the calibration families’ range, since logistic extrapolation can rank them as typical of calibration, as with the longest new proteins, and the gate’s descriptors omit length.

### What is only measured

Every comparison with baselines is empirical. Beyond the 0.01 margin, Prot-COVER’s advantage over group-weighted conformal prediction appears mainly under strong descriptor shift (S2 at *β* = 2), on one benchmark whose ten seeds resplit one pool of families, and, with ESM-1b but not ESM2, among new families without a detectable benchmark homolog (a secondary population); elsewhere it matches group weighting within the margin, and on proteins outside the benchmark no method reaches the target. ProtCOVER buys its smaller shortfall with larger sets under shift, and conditional calibration on one variant per family is within the margin of ProtCOVER in all but 1 of the 25 shifted cells.

### Scope and deployment

ProtCOVER is appropriate when a batch of unlabeled target families is available (a reference panel such as the unlabeled proteome could serve a single query, but the guarantee would then concern the panel’s shift); ProtCOVER-D can be used without one. Neither guarantees target coverage when predictor accuracy changes in ways the descriptors cannot capture. ProtCOVER-D’s shortfall was larger than ProtCOVER’s under descriptor shift but smaller on orphan families and, with ESM-1b, on the new proteins, and ProtCOVER’s default was chosen on the benchmark, which is development data. ProtCOVER outputs prediction sets, not ACMG/AMP evidence strengths; mapping sets to PP3/BP4 strengths is future work.

## 8 Conclusion

Calibrating a variant effect predictor variant by variant lets well-studied families set its error rate: on ProteinGym, Mondrian split conformal covers only 0.823–0.873 of pathogenic variants per family for all five predictors, without any shift. Treating families as the exchangeable unit removes this failure, and conditioning a family-weighted quantile regression on a cross-fitted density ratio guarantees coverage under family exchangeability and when the estimated ratio is correct. The empirical gain is in coverage and appears mainly under descriptor shift (S2); on orphan families and held-out embedding regions ProtCOVER matches group weighting. On proteins outside the benchmark, a pre-registered test with two predictors found variant-level calibration failing again and every method short of the target where the predictors were less accurate and the longest proteins lacked calibration support, neither of which the label-free ratio or the gate flags; conditioning narrowed the shortfall, within the margin of group weighting, without closing it. Descriptors that track a predictor’s accuracy without labels, and calibration support for proteins unlike any calibration protein, remain open.

## Code availability

Code and scripts that regenerate every number, table and figure from the public Prote-inGym release: https://github.com/wael-fahmy/protcover (MIT license).

