## Supplementary Appendix for "ProtCOVER: Family-Aware Conformal Prediction for Variant Effect Predictors on Unseen Protein Families"

#### A Proofs

Throughout, $F_k = N_k^{-1}\sum_{i=1}^{N_k}\delta_{S_{k,i}}$ is the score distribution of family $k$ (for the test family it involves unobserved labels and is used only in the analysis), and $\gamma_{K+1}(C) = F_{K+1}((-\infty,\hat{q}])$ because a test variant is covered iff its score is at most $\hat{q}$.

*Comparison step for* (2). Fix weights $p_1,\ldots,p_{K+1}$ and define the oracle measure $\mu = \sum_{k=1}^{K+1} p_k F_k$ and $q = \mathrm{Quant}_{1-\alpha}(\mu)$. The computable $\hat{\mu}$ moves the mass $p_{K+1}$ from $F_{K+1}$ to $+\infty$, so $\hat{\mu}((-\infty,u]) \le \mu((-\infty,u])$ for every $u$, hence $\hat{q} \ge q$ and $\gamma_{K+1}(C) \ge F_{K+1}((-\infty,q])$. Also $\mu((-\infty,q]) \ge 1-\alpha$ by right-continuity.

*Proof (Proposition 1).* With $\omega \equiv 1$, $p_k = 1/(K+1)$ and $q$ is a symmetric function of $(G_1,\ldots,G_{K+1})$. By exchangeability, $\mathbb{E}[F_{K+1}((-\infty,q])] = \mathbb{E}[(K+1)^{-1}\sum_k F_k((-\infty,q])] = \mathbb{E}[\mu((-\infty,q])] \ge 1-\alpha$, and the comparison step gives $\mathrm{Cov}(C) \ge 1-\alpha$.

*Proof (Theorem 1).* Let $L^o(b) = \sum_{k=1}^{K+1} N_k^{-1}\sum_i \rho_\tau(S_{k,i} - \Phi(D_k)^\top b)$ be the oracle objective, in which the test family enters with its true scores. It is convex, piecewise linear and nonnegative, so its minimizers form a nonempty closed convex polyhedron; let $\tilde{b}$ be the one of minimum Euclidean norm (a measurable function of the data, for example as the limit of the ridge-penalized minimizers as the penalty vanishes; only its symmetry is used), $\tilde{g} = \Phi^\top\tilde{b}$, and $\tilde{\gamma}_k = N_k^{-1}\sum_i \mathbf{1}\{S_{k,i} \le \tilde{g}(D_k)\}$.

*(a) First-order condition.* Let $f = \Phi^\top b_f$ with $f(D) \ge 0$ $P$-a.s.; then $f(D_k) \ge 0$ for every $k$ almost surely, which is all the argument uses. The right derivative of $t \mapsto \rho_\tau(S - \tilde{g}(D) - tf(D))$ at $t = 0$ is $f(D)(\mathbf{1}\{S \le \tilde{g}(D)\} - \tau)$ (if $S = \tilde{g}(D)$, the residual is $-tf(D) \le 0$ for $t > 0$). The right derivative of $t \mapsto L^o(\tilde{b} + tb_f)$ at 0 is therefore $\sum_{k=1}^{K+1} f(D_k)(\tilde{\gamma}_k - \tau)$, which is nonnegative at a minimizer.

*(b) Symmetry.* $L^o$, and hence $\tilde{b}$, is invariant to permutations of the families, so the pairs $(f(D_k),\tilde{\gamma}_k)$, $k \le K+1$, are exchangeable under $P^{K+1}$ and $\mathbb{E}_{P^{K+1}}[f(D_{K+1})(\tilde{\gamma}_{K+1} - \tau)] = (K+1)^{-1}\mathbb{E}\sum_k f(D_k)(\tilde{\gamma}_k - \tau) \ge 0$; the expectation exists because $\tilde{\gamma} \in [0,1]$ and $\mathbb{E}_P f < \infty$.

*(c) Change of measure.* For any bounded measurable $H$ of the families, $\mathbb{E}_{P^{K+1}}[f(D_{K+1})H] = \mathbb{E}_P f(D) \cdot \mathbb{E}_{P^K \times Q_f}[H]$. With $H = \tilde{\gamma}_{K+1} - \tau$, (b) gives $\mathbb{E}_{P^K \times Q_f}[\tilde{\gamma}_{K+1}] \ge \tau$.

*(d) The computable threshold dominates the oracle.* We show $\hat{q} \ge \tilde{g}(D_{K+1})$, which gives $\gamma_{K+1}(C) \ge \tilde{\gamma}_{K+1}$ and, with (c), the theorem. The argument follows Proposition 6 of [18] (imputing an upper bound $M$), with the supremum over the solution set in place of a min-norm solution, and $M \to \infty$. Write $\phi_t = \Phi(D_{K+1})$. If $L$ in (3) is unbounded below, $\hat{q} = +\infty$. Otherwise $A = \arg\min L$ is nonempty (a polyhedral function bounded below attains its minimum). Pick $M$ larger than every test score and than $\inf_{b\in A}\phi_t^\top b$, and let $L^M$ be $L^o$ with every test score replaced by $M$. Since $\rho_\tau(u) \ge \tau u$ with equality for $u \ge 0$, $L^M(b) = L(b) + \tau\phi_t^\top b + \rho_\tau(M - \phi_t^\top b) \ge L(b) + \tau M$, with equality when $\phi_t^\top b \le M$, which holds for some $b \in A$; hence $\min L^M = \min L + \tau M$ and $\arg\min L^M \subseteq A$. Let $b' \in \arg\min L^M$, $a = \tilde{g}(D_{K+1})$ and $a' = \phi_t^\top b'$. Adding $L^o(\tilde{b}) \le L^o(b')$ and $L^M(b') \le L^M(\tilde{b})$, the calibration terms cancel and $\sum_i [h_i(a) - h_i(a')] \le 0$, where $h_i(c) = \rho_\tau(S_{K+1,i} - c) - \rho_\tau(M - c)$ is nondecreasing in $c$ because $S_{K+1,i} \le M$ and $\rho_\tau$ is convex. If $a' < a$, then $h_i(a') \le h_i(a)$ for every $i$, so both optimality inequalities hold with equality and $\tilde{b} \in \arg\min L^M \subseteq A$. In either case $\hat{q} = \sup_{b\in A}\phi_t^\top b \ge a$.

The case $f \equiv 1$ gives the statement under A1 (for exchangeable rather than i.i.d. families, (b) applies directly with $f \equiv 1$).

*Computing the supremum.* Let $L'$ be $L$ with the test term's weight raised from $\tau$ to $\tau' > \tau$, so $L'(b) = L(b) - (\tau' - \tau)\phi_t^\top b$. If $b \in \arg\min L$ and $b' \in \arg\min L'$, adding $L(b) \le L(b')$ and $L'(b') \le L'(b)$ gives $(\tau' - \tau)\,\phi_t^\top(b' - b) \ge 0$, so every minimizer of $L'$ has $\phi_t^\top b' \ge \hat{q}$. If $L$ is unbounded below along a direction $d$, its recession slope along $d$ is negative, which forces $\phi_t^\top d > 0$, so $L'$ is unbounded along $d$ as well. If $L$ is bounded but $\hat{q} = +\infty$, then $A$ contains points with $\phi_t^\top b \to \infty$ and $L'$ is unbounded below on $A$. In every case every exact minimizer of $L'$ with $\tau' = \tau + 10^{-4}$ has fitted value at least $\hat{q}$, and an unbounded $L'$ gives $+\infty$; these fitted values all equal $\hat{q}$ when $\tau' - \tau$ is small enough for the data (the optimal value is concave and piecewise linear in $\tau'$, with finitely many breakpoints, and its right derivative at $\tau$ is $-\hat{q}$), and is otherwise conservative. A floating-point solver finds a minimizer of $L'$ only up to its tolerances, and near a tie of $L$ it can return the other end of the tie, below $\hat{q}$, so every computed threshold is certified. For the basis that the solver returns, $b$ solves the $p$ interpolation equations of its basic scores exactly; every other score's dual variable is set to the bound that the sign of its exact residual dictates; and $b$ minimizes $L'$ if the basic dual variables, which then solve a $p \times p$ system, lie within their bounds (a sufficient condition: with scores on the fitted surface it can fail for an optimal $b$, a case the exact simplex below resolves). This check is done in exact rational arithmetic, with $\tau = 1 - \alpha$, $\tau'$ and the weights $1/N_k$ exact (floating-point evaluation with rigorous error bounds decides almost every case), and the certified fitted value is rounded up. A basis can pass the solver's tolerances and fail the check: a score whose residual is below those tolerances may belong at its other bound. If neither the returned basis nor that of a cold re-solve with tight tolerances passes, a bounded dual simplex in exact rational arithmetic takes over, started from the solver's basis with every nonbasic dual variable at the bound its exact residual sign dictates (a dual-feasible start; bound-flipping ratio test, Bland's smallest-index rule on degenerate steps). It ends with an exactly optimal basis, whose optimality it checks again, or with an exact certificate that $L'$ is unbounded; the solver's unboundedness verdicts are checked the same way. Every computed threshold is therefore the exact optimal fitted value of $L'$, rounded up, except that a pivot cap, never reached, would set it to $+\infty$. By Lemma 2 the computed sets keep every guarantee for the stored double-precision scores and features. Standardizing the descriptors (ProtCOVER-D) is an invertible affine reparametrization of $b$ that does not change $\mathcal{F}$, so fitted values, and $\hat{q}$, are unchanged. With $\Phi \equiv 1$, (3) gives the HCP threshold (2), except that when the pooled distribution $\hat{\mu}$ reaches $1 - \alpha$ exactly at that threshold it takes the next pooled calibration score, or $+\infty$ if there is none.

*Proof (Corollary 1).* Let $\phi(g) \in [0, 1]$ be the expected per-family coverage of a test family $g$ over $G_1, \ldots, G_K \sim P$. For $f \in \mathcal{F}$, $f \geq 0$, Theorem 1 gives $\mathbb{E}_{Q_f}\phi \geq 1 - \alpha$, and $|\mathbb{E}_Q\phi - \mathbb{E}_{Q_f}\phi| \leq d_{\mathrm{TV}}(Q, Q_f)$. For $\Phi = (1, \hat{w})$, condition on $\hat{w}$: $f \equiv 1$ and $f = \hat{w}$ are nonnegative members of $\mathcal{F}$ with $Q_1 = P$ and $Q_{\hat{w}} = \hat{Q}$.

*Proof (Proposition 2).* Under A2 the joint law of $(G_1, \ldots, G_{K+1})$ has density $\prod_{k \leq K} \pi(g_k) \cdot w(d_{K+1})\pi(g_{K+1})$ with respect to a product base measure, where $\pi$ is a density of $P$, so the sequence is weighted exchangeable with weight functions $1, \ldots, 1, w$ [41, Def. 1, Lemma 2]. Let $E$ be the event that the unordered collection $\{G_1, \ldots, G_{K+1}\}$ equals $\{g_1, \ldots, g_{K+1}\}$. As in the proof of Lemma 3 of [41], $\mathbb{P}(G_{K+1} = g_k \mid E) = w(d_k)/\sum_j w(d_j) = p_k$ (summing over coincident values). The $p_k$, $\mu$ and $q$ are functions of $E$, so $\mathbb{E}[F_{K+1}((-\infty, q]) \mid E] = \mu((-\infty, q]) \geq 1 - \alpha$, and the comparison step gives the first claim. For $\hat{w}$ independent of the families, the procedure is invariant to rescaling the weights, so conditionally on $\hat{w}$ it is the procedure of the first claim for the shift $\hat{Q}$; the total-variation argument of Corollary 1 gives the second claim.

*Proof (Lemma 1).* The ratio for fold $j$ is fitted on the fitting calibration families and the target families outside fold $j$, rescaled to mean one over the fitting calibration families and clipped by a fixed rule; no other family enters it. If all families are mutually independent (as under A2, and in Corollary 1 for any test law $Q$), $\hat{w}_j$ is independent of the calibration families used in (2) and (3) and of each family of fold $j$, which, conditionally on $\hat{w}_j$, are still i.i.d. $P$ and $Q$. Each query family of fold $j$ plays the role of $G_{K+1}$ in Corollary 1 and Proposition 2. If the non-training families are exchangeable (a random split of an exchangeable pool, given the training families), then conditionally on the fitting calibration families and the target families outside fold $j$, each in their given order (the classifier depends on which families are of which class and, through its cross-validation folds, on their order), $\hat{w}_j$, hence $\Phi$, is fixed, and the other calibration families together with any one family of fold $j$ are exchangeable; step (b) of the proof of Theorem 1 with $f \equiv 1$ applies directly to them, and (d) is unchanged. A unit test in the released code checks that a family's weight does not change when the other calibration families change.

*Proof (Proposition 3).* Fix $y$ and let $\mathcal{I}_y = \{k \leq K+1 : N_{k,y} > 0\}$. Conditionally on $\mathcal{I}_y$, with $K+1 \in \mathcal{I}_y$, the families indexed by $\mathcal{I}_y$ are exchangeable under A1 (the event $\{\mathcal{I}_y = I\}$ is invariant under permutations of $I$) and independent in the other settings, with laws $P_y = P(\cdot \mid N_y > 0)$ for calibration families and the test law conditioned on $N_y > 0$. Because $\mathbf{1}\{N_y > 0\}$ is a function of $G$, a test law with density $f(D)/\mathbb{E}_P f$ with respect to $P$ has, after conditioning, density proportional to $f(D)$ with respect to $P_y$. The label-$y$ procedures are the procedures above applied to these families with $F_k$ replaced by the label-$y$ score distribution $F_k^y$ (weights $1/N_{k,y}$), so every proof applies verbatim, and the normalizing constants cancel.

*Proof (Lemma 2).* $\mathbf{1}\{Y \in C'(X)\} \geq \mathbf{1}\{Y \in C(X)\}$ pointwise, so every average and expectation of the indicators is at least as large for $C'$.

### B Protocol Details

*Predictors.* The five unsupervised predictors are all the non-clinical models in ProteinGym v1.3's clinical score files (columns ESM1b, EVE, GEMME, TranceptEVE_L, PoET; PoET is the 200M model). The other columns are clinical predictors trained or tuned on clinical labels.

*Families and homology.* MMseqs2 release 18-8cc5c. Clusters: `cluster` with minimum identity 0.3 (0.5 for the sensitivity check), coverage 0.8 and coverage mode 0 (both sequences). Homology search: all against all, sensitivity 7.5, $E \leq 10$, at most 500 hits per query; hits are then filtered at $E \leq 10^{-3}$. Homology components additionally require identity $\geq 0.3$ and coverage $\geq 0.5$ of both sequences; the strictest definition takes connected components of all hits with $E \leq 10^{-3}$ (1,024 families, 778 single proteins, one component of 867 proteins). That component holds 45% of the benchmark's variants and, across the ten S1 draws, falls in training three times, in calibration four times and in the test set three times; family-level calibration and family-weighted coverage give it the weight of one family, whereas it dominates variant-level calibration whenever it calibrates and variant-weighted metrics whenever it is tested.

*Descriptors.* Per protein: maximum identity to any training protein over hits with $E \leq 10^{-3}$ (0 without a hit); $\log(1 + N_{\text{eff}})$ with $N_{\text{eff}}$ the sum of ProteinGym's MSA sequence weights; $\log(N_{\text{eff}}/L + 10^{-3})$ with $L$ the alignment length (written $\log N_{\text{eff}}$ per position); cosine distance of the mean-pooled last-layer (layer 12) embedding of `esm2_t12_35M_UR50D` to the nearest training protein (sequences over 1022 residues are embedded in windows of 1022 with stride 511 and residue representations averaged over windows); log length. Per family: the maximum of maximum identity and the minimum of embedding distance over member proteins, the mean of the other protein descriptors, and, in the transductive run only, log of the family's number of scored variants. Descriptors are recomputed for every split because they depend on its training families.

*Splits.* S2 uses $z$, the mean of the standardized maximum identity and $\log(1 + N_{\text{eff}})$ over non-training families, and Efraimidis–Spirakis keys $\log(u)\, e^{\beta z}$ with uniforms $u$ shared across $\beta$, so the sweep is coupled and $\beta = 0$ is S1. S3 draws test families uniformly from orphan families and training and calibration families uniformly from the rest. S4 regions are fixed once, before any S4 run: each family's embedding is the mean of its members' unit-normalized ESM2 embeddings, renormalized; after centering, the first 50 principal components are clustered by $k$-means ($k = 20$, ten restarts, seed 0). Each seed permutes the regions and adds whole regions to the test set until it holds 30% of families (the last region is split at random); training and calibration families are drawn uniformly from the remaining families. S4s permutes the regions in the same way but adds a region only if the test set stays within 30% of families, skipping the others, so that every test region is held out whole (590–624 test families per seed); training and calibration families are drawn uniformly from the remaining families in the proportions 40:30. S5 keeps the S1 training families; $z$ is the standardized mean over the five predictors of each family's within-family AUC (families without both classes get the median), and test families are drawn as in S2 with $\beta = 1$. The ten seeds resplit one fixed pool of families, so they are not independent samples of the population of families.

*Base model and conformal methods.* The base model standardizes its feature (the raw score clipped to the training 0.1% and 99.9% quantiles; in the transductive run also the score standardized over the protein's scored variants) and fits $L_2$-regularized logistic regression ($C = 1$). Earlier versions of this work (in the public repository) used the transductive base model as the primary configuration; it is now the deployment-realistic one, as in both external evaluations. Dunn et al.'s repeated subsampling uses 50 subsamples. GWCP uses nine bins: maximum identity zero, at most the median of the positive values, or above it, crossed with $\log(1+N_{\text{eff}})$ terciles over calibration and test families; its target proportions are the bin frequencies over all test families (label-free), and it pools at the family level like HCP. For $\hat{w}$, one third of the calibration families fit the classifier (logistic regression with ten candidate penalties, five-fold cross-validated log loss) and the test families are split into five folds. ProtCOVER uses the features $(1, \hat{w}_j)$ on the remaining calibration families for queries of fold $j$; ProtCOVER-D uses an intercept and maximum identity to training, $\log(1 + N_{\text{eff}})$, $\log N_{\text{eff}}$ per position and embedding distance, standardized over all calibration families. Both feature classes were fixed in the configuration before the conditional methods were first run. They are finite-dimensional and linear because for such classes the guarantee needs no approximation; regularized or kernel classes [18] would hold only up to a regularization term, and we do not study them. The alignment-free run removes $\log(1 + N_{\text{eff}})$ and $\log N_{\text{eff}}$ per position from the density ratio, from ProtCOVER-D's features and from the gate, and GWCP crosses maximum identity with embedding-distance terciles instead. The linear programs are solved with HiGHS 1.11 (`highspy`), warm-started across test families, with a cold re-solve and the exact simplex of Appendix A as fallbacks.

*Threshold audit.* Across the 14 runs that use conditional calibration, 17,866,020 thresholds were computed: 17,821,146 were certified from the solver's basis, 16,767 after a cold re-solve and 1,186 by the exact simplex, and 26,921 are infinite, each confirmed exactly; no unboundedness verdict of the solver was overturned, and no computation reached the pivot cap. Sets include a label when its score is at most the threshold plus $10^{-9}$, a guard against rounding of the stored scores that can only enlarge sets (Lemma 2). As an independent check of the implementation, `scripts/audit_thresholds.py` recomputes 1,800 of them independently (ProtCOVER and ProtCOVER-D; seed 0; S1, S2 at $\beta \in \{0.5, 1, 2\}$, S3 and S4; all five predictors; both classes; three random test families per fold): on the primal program with the exact test weight $\tau$, it minimizes $L$ and then maximizes the query's fitted value subject to $L(b) \leq L^\star$, both with tolerances of $10^{-10}$. Every reference computation succeeded, 25 of them only after rounding forced a relative slack of $10^{-12}$ on $L^\star$. The two computations agree on whether the threshold is infinite (both are, in 4 cases). The production threshold exceeds the reference by more than $10^{-6}$ in 7 cases, by at most $1.5 \times 10^{-1}$; there $\tau + 10^{-4}$ crosses a breakpoint of the optimal value, because with $\tau + 10^{-6}$ the same thresholds match the reference to $4.4 \times 10^{-7}$ (with $\tau + 10^{-5}$, all but one of them). It is below the reference in 23 cases, by at most $6.8 \times 10^{-8}$: 22 of them needed slack, which can raise the reference's fitted value, and the remaining one is below by $1.9 \times 10^{-9}$, within the reference's tolerances, so all are within the precision of the check. Elsewhere the two agree to $1.6 \times 10^{-9}$. No audited test variant changes membership between the two thresholds (0 lost, 0 added).

*External test set.* ClinVar `variant_summary` (October 2026) and `hgvs4variation` map VariationIDs to protein changes in the benchmark's RefSeq proteins; we keep single-nucleotide missense variants on GRCh38 with at least one review star, labeled pathogenic/likely pathogenic or benign/likely benign, whose VariationID is absent from the June 2023 release and whose protein change is not in the benchmark. Excluded as a precaution: VariationIDs present in the July 2023 release and changes that were in the June 2023 release under another VariationID. ESM-1b scores are the precomputed log-likelihood ratios of Brandes et al. [9], oriented as in the benchmark; for proteins of at most 1,022 residues they equal the benchmark's to rounding, so only those are used (longer proteins hold 43% of the new variants). The build script, the pre-registration and its amendment, and the outcome are in the repository (`docs/external_results.md`). The pre-registration was written after a sanity check of ESM-1b's AUC on these data, and the evaluation was rerun, with identical results, after each of the two fixes to the threshold computation (certification, then the exact simplex). All four hypotheses were supported in the primary and the secondary analysis: Mondrian split under-covers pathogenic variants per family in S1 (H1; 0.873 [0.864, 0.883]), HCP covers both classes at 0.89 or more (H2; 0.939 and 0.918), ProtCOVER-D's shortfall over the shifted settings is at most HCP's and GWCP's (H3; ProtCOVER was added to H3 after development had singled it out, though before the run), and abstention does not lower the error of retained singleton sets (H4, whose decision rule we fixed after the run, so it is exploratory). ESM-1b separates these new variants better than the benchmark variants of the same proteins (per-protein AUC 0.930 against 0.899), as expected if recent classifications used computational evidence (PP3/BP4), a caveat stated in the pre-registration. The higher AUC is not confined to the classifications where such evidence weighs most (exploratory, among proteins with both labels): it is 0.931 for "likely" classifications (834 proteins) and 0.934 for definitive ones (103), and 0.931 for single-submitter variants against 0.962 with several submitters or expert review (45), so these data cannot tell how much computational evidence contributes. An exploratory rerun, not pre-registered, keeps proteins of any length and scores the benchmark variants with Brandes et al.'s values as well: the external test sets grow from 4,050 to 7,117 variants on average, and no verdict changes (Mondrian split 0.870 in S1; over the shifted settings HCP falls short by 0.006 and no shift-aware method falls short).

*New-protein test set.* The same ClinVar files and variant filters, without a date filter, in approved HGNC protein-coding genes outside the benchmark (excluded by gene symbol, GeneID and UniProt accession). Each gene contributes the canonical UniProt entry that Brandes et al. scored, reached through a RefSeq protein with the same numbering, and the wild type is checked at every position. Families are MMseqs2 clusters of the new proteins with the benchmark's parameters; every new protein is also searched against every benchmark protein, which gives its maximum identity to any training set and defines the secondary population (no hit with $E \leq 10^{-3}$). Benchmark variants without a Brandes et al. score (2,755) are dropped from training and calibration, so that one score function covers both sides at any length. Clustering the new proteins jointly with the benchmark would put 1,581 of them (656 families) into a family with a benchmark protein; as pre-registered they stay in the primary population, and the secondary population excludes every family with a detectable benchmark homolog. The second predictor, ESM2 (`esm2_t30_150M_UR50D`), is scored by the wild-type marginal log-likelihood ratio [31], one forward pass per window of 1,022 residues with stride 511 and log-probabilities averaged over the windows that cover a position, negated so that higher means more pathogenic, on the same variants. Before the evaluation, as pre-registered, its per-protein mean AUC on the benchmark variants that train and calibrate it was checked: 0.808 over 2,423 proteins (ESM-1b 0.892), above the 0.5 that would have meant an orientation or position bug. The first ESM-1b run used the code before threshold certification; it was rerun after that fix and again after the exact simplex, with identical results, and the first run's outputs are in the repository with the build script and both pre-registrations.

*Gate.* The gate acts on the family of a query protein, so all variants of that family share one decision. With standardized support descriptors $\tilde{d}$, the kernel weights $\kappa_{tk} = \exp(-\|\tilde{d}_k - \tilde{d}_t\|^2/2b^2)$ give the effective number of calibration families that resemble query family $t$, $\mathrm{ESS}_t = (\sum_k \kappa_{tk})^2/\sum_k \kappa_{tk}^2$, and the test-point mass $\pi_t = p_{K+1}$ of ProtCOVER-W measures how much of the weighted quantile's mass sits on the query itself. The gate abstains when $\mathrm{ESS}_t < m$ or $\pi_t > \pi_{\max}$: the first catches queries with few similar calibration families, the second queries that the ratio marks as far more typical of the targets than of the calibration families. Tuned under the 10% budget (a cap), it abstained on at most 1.4% of test families except at $\beta = 2$ (0.1%–5.2%).

*Gate tuning.* An inner split of each seed's training families holds out 40% as inner training families (descriptors are recomputed relative to them) and draws 30% as inner test families with the S2 sampler at $\beta = 1$; the rest are inner calibration families. Grids: kernel scale $\{0.1, 0.2, 0.35, 0.5, 1\}$ times the median pairwise distance of standardized support descriptors, $m \in \{0, 2, 5, 10, 20, 40\}$, $\pi_{\max} \in \{1, 0.05, 0.02, 0.01, 0.005\}$, abstention budget 10%, coverage tolerance 0.005.

*Metrics.* Coverage metrics average per-family coverage over families; set size and singleton rate average over variants. The under-covered fraction uses overall per-family coverage (both classes) for families with at least ten variants, for marginal and class-conditional methods alike; its binomial reference is the expected fraction under $\mathrm{Binomial}(N_k, 1-\alpha)$. Bootstrap intervals use 1000 resamples of test families within each seed, holding that seed's calibration draw fixed, so they are conditional on the splits; seed-to-seed variation is reported separately. Random abstention in risk-coverage curves is averaged over 100 random family orders per cell. The 0.01 margin was pre-registered for the external evaluations; its placement between two noise levels of the benchmark (Section 6.2) was computed afterwards, on development data. With half or twice the margin, ProtCOVER's shortfall is smaller than GWCP's in 11 and 3 of the 25 shifted cells and larger in 0 and 0 (against HCP: 19 and 7, larger in 0 and 0). The 25 cells and three comparisons carry no multiplicity correction, and seeds resplit one pool, so these counts describe this benchmark rather than test hypotheses.

#### C Synthetic Validation

We simulate families with a descriptor $d \sim N(0, 1)$, random offsets, and slopes on the score whose mean depends on $d$, with a fixed base model, so that the truth is known (300 replicates, Monte Carlo standard error of every coverage at most 0.002). Under A1 with larger families being easier, variant-level split conformal has variant-weighted coverage 0.900 but family-weighted coverage 0.831, while HCP attains 0.906 and single subsampling 0.902. Under exact family-level covariate shift (A2, test descriptors $N(-\beta, 1)$), HCP drops to 0.826 at $\beta = 1$ and 0.734 at $\beta = 2$; weighting with the true ratio attains 0.911 and 0.950, and with cross-fitted estimated ratios 0.916 and 0.951 (class-conditional pathogenic: 0.916 and 0.947). The excess at $\beta = 2$ is the price of the test-point mass, which grows when few calibration families resemble the test family.

For the class-conditional versions we report the worse of the two class-wise coverages (target 0.90). Under the Gaussian shift at $\beta = 1$ and $\beta = 2$, it is 0.818 and 0.714 for HCP; conditional calibration on $(1, w)$ with the true ratio, the case Theorem 1 covers exactly, attains 0.909 and 0.922; ProtCOVER, with the cross-fitted ratio, 0.913 and 0.939; ProtCOVER-D, whose features $(1, d)$ do not span this ratio, 0.906 and 0.921; and reweighting with the true ratio 0.910 and 0.948. To test the theorem where the descriptor features are exact, a second shift draws calibration descriptors from $U(-2, 2)$ and test descriptors with density $(2 - d)/8$ on $[-2, 2]$, so that $w(d) = (2 - d)/2$ is affine in $d$: HCP attains 0.859, and ProtCOVER-D 0.905, conditional calibration on the true ratio 0.905, ProtCOVER 0.906 and reweighting with the true ratio 0.904. The central checks are also unit tests in the released code.

### D Additional Results

*Robustness runs.* With the transductive base model, Mondrian split still under-covers a class per family in S1 for every predictor (worse class 0.749–0.837; for TranceptEVE the benign class), and the mean shortfall under shift is 0.001 for ProtCOVER, 0.007 for GWCP and 0.023 for HCP. Without alignment depth, GWCP falls short as much as HCP (both 0.019) and ProtCOVER by 0.011. In ablations, reweighting variants instead of families leaves a mean worse-class shortfall of 0.054 over the shifted cells (pathogenic coverage in Table 4). The strictest family definition leaves no test protein with a detectable homolog, even through a shared domain, among the training or calibration proteins.

*Pooling, feature classes, information and the ratio's share.* Over the shifted cells, conditional calibration on one random variant per family and class falls short by 0.002 on average (worst cell 0.012), against 0.001 (0.008) for ProtCOVER, which pools every variant. ProtCOVER's worse-class coverage is higher with a paired interval above zero in 12 cells and lower in 2, and its pathogenic coverage varies less across seeds (s.d. 0.016 against 0.020); at matched coverage both need the same set size (1.267 against 1.268), and beyond the margin pooling is better in 1 of the 25 cells and worse in 0. Adding the descriptors to $(1, \hat{w})$ lowers the shortfall to 0.000 (worst 0.002) at a larger set size (1.316 against 1.294), and quadratic descriptor features lower ProtCOVER-D's shortfall from 0.004 to 0.001. Two comparisons separate the method from the information it uses. GWCP whose groups are the ninths of ProtCOVER's ratio (for each test fold, over the calibrating families and the fold's test families, with target proportions from the fold's test families) falls short by 0.006 (worst 0.019), against 0.007 for GWCP on its nine descriptor bins; conditional calibration on GWCP's information, a constant and the indicators of its bins (every function constant on the bins), falls short by 0.000 (worst cell 0.000) but abstains on up to 13.4% of a cell's test families (ProtCOVER: 0.3%), in the 5 cells of S2 at $\beta = 2$, because by the boundedness condition of Section 4 a bin needs at least $\tau/(1 - \tau) = 9$ calibration families with variants of a class for a finite threshold at $\alpha = 0.1$; on the families it does not abstain on, its worse-class coverage in the worst cell is 0.900, so abstention alone does not close the gap. With ProtCOVER's information, GWCP comes no closer to ProtCOVER, whose shortfall is still smaller beyond the margin in 4 cells (as many as against GWCP with its own bins) and larger in none, while with GWCP's information conditional calibration falls short less than GWCP: the gain comes mainly from the method, not from the ratio's information, and conditioning on the single feature $\hat{w}$ rather than on nine bin indicators keeps abstention rare. Fitting the ratio on 20% or 50% of the calibration families instead of a third gives ProtCOVER a shortfall of 0.001 and 0.000, within the margin of the default in every cell (Table 6).

Tables 7–10 give paired differences between ProtCOVER and HCP, GWCP and ProtCOVER-W (class-conditional versions, $\alpha = 0.1$, mean over seeds) with 95% paired family-bootstrap intervals, which are conditional on the splits. Singleton error is the fraction of single-label sets that exclude the true label.

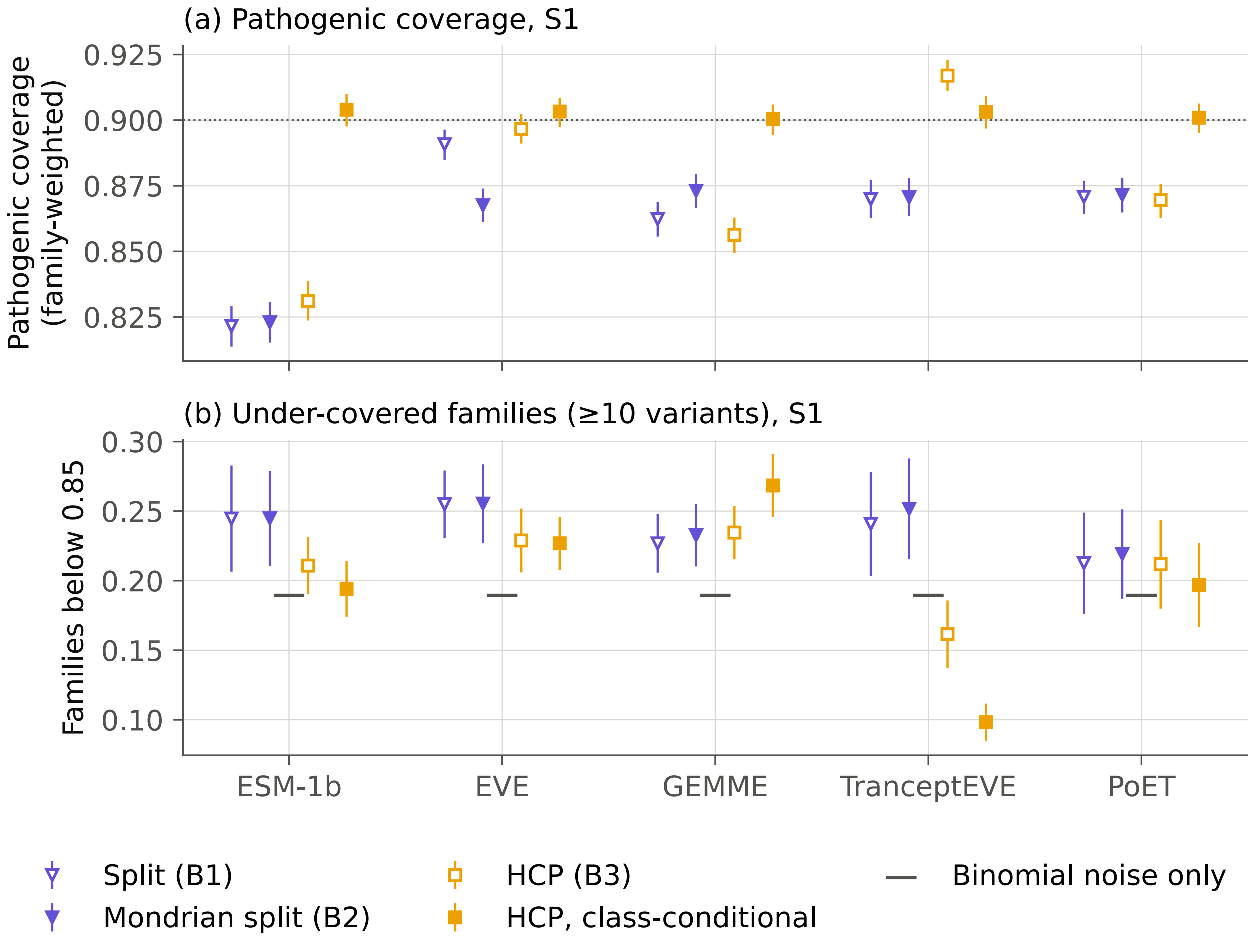


**Fig. 1.** Exchangeable families (S1, $\alpha = 0.1$). (a) Family-weighted pathogenic coverage $\mathrm{Cov}_1$ with 95% family-bootstrap intervals; the dotted line is the target. (b) Fraction of families with at least ten variants whose coverage is below 0.85 (mean and s.d. over seeds); the gray dash is the expected fraction if every family had coverage exactly 0.90.

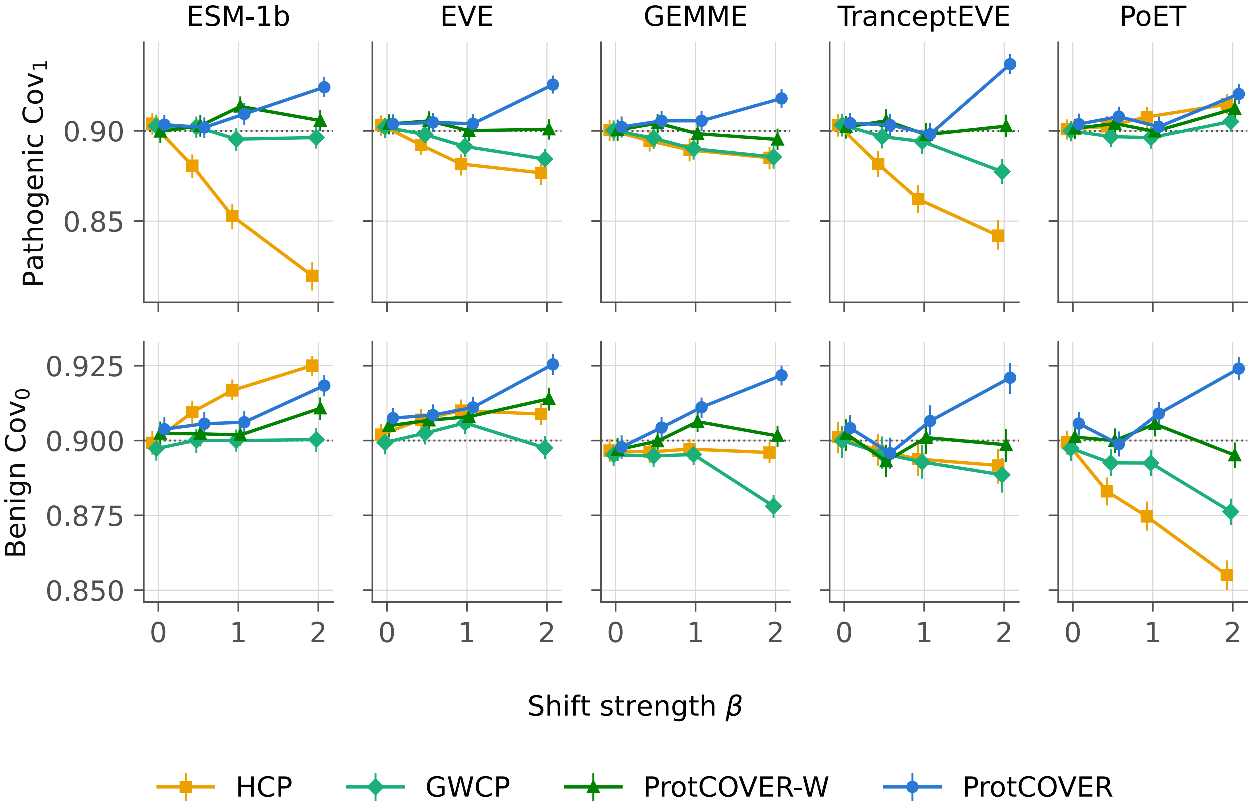


**Fig. 2.** Class-conditional coverage of both classes under increasing descriptor shift (S2; $\beta = 0$ is S1), $\alpha = 0.1$, with 95% family-bootstrap intervals; methods are offset slightly in $\beta$. Mondrian split conformal, far below target, is omitted (Fig. 1).

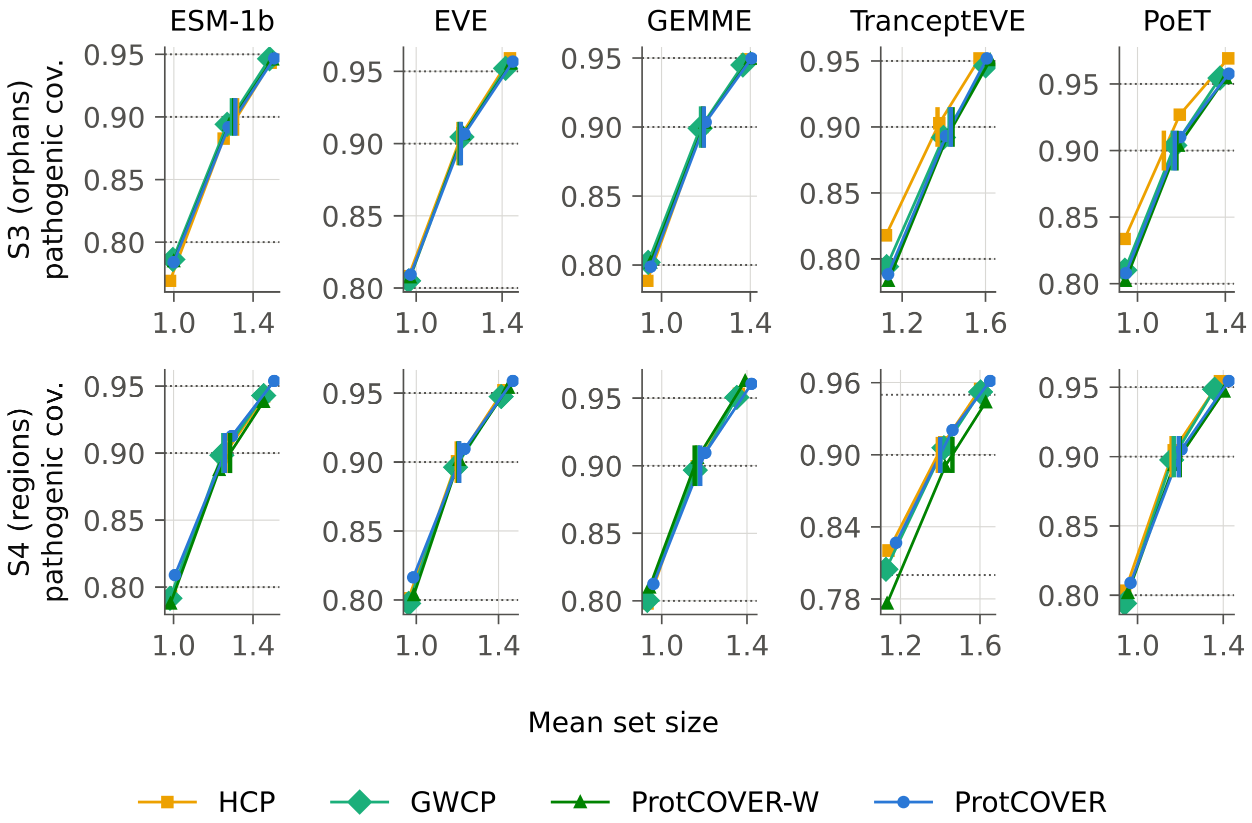


**Fig. 3.** Pathogenic coverage against mean set size for $\alpha \in \{0.2, 0.1, 0.05\}$ (left to right along each curve), class-conditional methods, on the two shifts defined without the descriptors. Dotted lines mark the three targets. A short vertical bar on the 0.90 line marks where a method's curve crosses pathogenic coverage 0.90, found by linear interpolation between its two neighboring levels: the set size the method needs for that coverage, which the last row of Table 2 averages over all shifted cells, S2 included. The methods lie close to one coverage-size curve, so a method that covers more also returns larger sets; the exceptions are HCP, above the curve for TranceptEVE and PoET in S3, and ProtCOVER-W, below it for TranceptEVE in S4.

**Table 3.** Family-weighted pathogenic / benign coverage ($\mathrm{Cov}_1$ / $\mathrm{Cov}_0$) of class-conditional methods at $\alpha = 0.1$ (target 0.90), per predictor, mean over ten seeds.

| Method | ESM-1b | EVE | GEMME | TranceptEVE | PoET |
|---|---|---|---|---|---|
| *S1 (exchangeable)* | | | | | |
| Mondrian split (B2) | 0.823 / 0.914 | 0.868 / 0.914 | 0.873 / 0.921 | 0.871 / 0.802 | 0.872 / 0.906 |
| HCP (B3) | 0.904 / 0.899 | 0.903 / 0.902 | 0.900 / 0.897 | 0.903 / 0.901 | 0.901 / 0.899 |
| GWCP (B4) | 0.903 / 0.897 | 0.902 / 0.899 | 0.900 / 0.895 | 0.903 / 0.900 | 0.900 / 0.897 |
| ProtCOVER-W | 0.900 / 0.902 | 0.904 / 0.905 | 0.900 / 0.897 | 0.902 / 0.902 | 0.901 / 0.901 |
| ProtCOVER-D | 0.910 / 0.906 | 0.907 / 0.908 | 0.906 / 0.902 | 0.910 / 0.906 | 0.909 / 0.902 |
| **ProtCOVER** | 0.903 / 0.904 | 0.904 / 0.908 | 0.902 / 0.898 | 0.904 / 0.904 | 0.904 / 0.906 |
| *S2*, $\beta = 0.5$ | | | | | |
| Mondrian split (B2) | 0.792 / 0.924 | 0.852 / 0.918 | 0.868 / 0.919 | 0.851 / 0.808 | 0.876 / 0.895 |
| HCP (B3) | 0.881 / 0.910 | 0.892 / 0.907 | 0.894 / 0.896 | 0.882 / 0.897 | 0.902 / 0.883 |
| GWCP (B4) | 0.902 / 0.900 | 0.898 / 0.902 | 0.896 / 0.895 | 0.897 / 0.896 | 0.897 / 0.893 |
| ProtCOVER-W | 0.903 / 0.902 | 0.905 / 0.907 | 0.904 / 0.900 | 0.906 / 0.893 | 0.904 / 0.900 |
| ProtCOVER-D | 0.909 / 0.906 | 0.905 / 0.908 | 0.902 / 0.904 | 0.904 / 0.904 | 0.905 / 0.900 |
| **ProtCOVER** | 0.902 / 0.906 | 0.905 / 0.909 | 0.906 / 0.904 | 0.903 / 0.896 | 0.908 / 0.899 |
| *S2*, $\beta = 1$ | | | | | |
| Mondrian split (B2) | 0.767 / 0.931 | 0.843 / 0.921 | 0.863 / 0.919 | 0.838 / 0.811 | 0.880 / 0.888 |
| HCP (B3) | 0.853 / 0.917 | 0.882 / 0.910 | 0.889 / 0.897 | 0.862 / 0.894 | 0.908 / 0.875 |
| GWCP (B4) | 0.895 / 0.900 | 0.891 / 0.906 | 0.890 / 0.895 | 0.894 / 0.893 | 0.896 / 0.892 |
| ProtCOVER-W | 0.913 / 0.902 | 0.900 / 0.908 | 0.898 / 0.906 | 0.898 / 0.901 | 0.900 / 0.906 |
| ProtCOVER-D | 0.907 / 0.910 | 0.896 / 0.912 | 0.893 / 0.906 | 0.895 / 0.907 | 0.907 / 0.901 |
| **ProtCOVER** | 0.909 / 0.906 | 0.904 / 0.911 | 0.906 / 0.911 | 0.898 / 0.907 | 0.902 / 0.909 |
| *S2*, $\beta = 2$ | | | | | |
| Mondrian split (B2) | 0.744 / 0.939 | 0.835 / 0.922 | 0.862 / 0.916 | 0.833 / 0.813 | 0.889 / 0.874 |
| HCP (B3) | 0.820 / 0.925 | 0.877 / 0.909 | 0.885 / 0.896 | 0.842 / 0.892 | 0.915 / 0.855 |
| GWCP (B4) | 0.896 / 0.900 | 0.884 / 0.898 | 0.885 / 0.878 | 0.877 / 0.889 | 0.905 / 0.876 |
| ProtCOVER-W | 0.906 / 0.911 | 0.901 / 0.914 | 0.895 / 0.902 | 0.903 / 0.899 | 0.912 / 0.895 |
| ProtCOVER-D | 0.907 / 0.918 | 0.892 / 0.908 | 0.885 / 0.906 | 0.884 / 0.908 | 0.917 / 0.886 |
| **ProtCOVER** | 0.924 / 0.918 | 0.926 / 0.925 | 0.918 / 0.922 | 0.937 / 0.921 | 0.920 / 0.924 |
| *S3 (orphan families)* | | | | | |
| Mondrian split (B2) | 0.785 / 0.926 | 0.867 / 0.911 | 0.866 / 0.933 | 0.883 / 0.777 | 0.900 / 0.899 |
| HCP (B3) | 0.883 / 0.911 | 0.907 / 0.903 | 0.900 / 0.911 | 0.903 / 0.886 | 0.927 / 0.890 |
| GWCP (B4) | 0.894 / 0.906 | 0.905 / 0.904 | 0.899 / 0.905 | 0.892 / 0.909 | 0.904 / 0.904 |
| ProtCOVER-W | 0.894 / 0.912 | 0.907 / 0.907 | 0.903 / 0.913 | 0.893 / 0.919 | 0.904 / 0.914 |
| ProtCOVER-D | 0.900 / 0.912 | 0.907 / 0.907 | 0.904 / 0.913 | 0.897 / 0.914 | 0.914 / 0.908 |
| **ProtCOVER** | 0.892 / 0.913 | 0.907 / 0.908 | 0.904 / 0.915 | 0.893 / 0.914 | 0.910 / 0.910 |
| *S4 (held-out regions)* | | | | | |
| Mondrian split (B2) | 0.810 / 0.920 | 0.862 / 0.911 | 0.877 / 0.928 | 0.896 / 0.782 | 0.876 / 0.908 |
| HCP (B3) | 0.897 / 0.902 | 0.901 / 0.901 | 0.900 / 0.904 | 0.910 / 0.887 | 0.904 / 0.902 |
| GWCP (B4) | 0.898 / 0.902 | 0.896 / 0.900 | 0.897 / 0.901 | 0.906 / 0.894 | 0.898 / 0.906 |
| ProtCOVER-W | 0.887 / 0.905 | 0.902 / 0.899 | 0.906 / 0.895 | 0.890 / 0.915 | 0.894 / 0.913 |
| ProtCOVER-D | 0.890 / 0.914 | 0.909 / 0.905 | 0.911 / 0.907 | 0.889 / 0.904 | 0.906 / 0.915 |
| **ProtCOVER** | 0.913 / 0.905 | 0.910 / 0.904 | 0.910 / 0.909 | 0.920 / 0.906 | 0.905 / 0.917 |

**Table 4.** Ablations of ProtCOVER-W, the reweighting variant: pathogenic coverage $\mathrm{Cov}_1$ at $\alpha = 0.1$, mean (minimum) over the five predictors, class-conditional versions. The clipping level $c = 20$ dates from its first version, before the conditional methods existed; ProtCOVER's own descriptor set and clipping are not ablated.

| Configuration | S2, $\beta = 1$ | S2, $\beta = 2$ | S3 | S4 |
|---|---|---|---|---|
| *Components* | | | | |
| (i) family-level calibration (HCP) | 0.879 (0.853) | 0.868 (0.820) | 0.904 (0.883) | 0.902 (0.897) |
| (i)+(ii), conditional on $\hat{w}$ (ProtCOVER) | 0.904 (0.898) | 0.925 (0.918) | 0.901 (0.892) | 0.912 (0.905) |
| (i)+(ii), conditional on descriptors (ProtCOVER-D) | 0.899 (0.893) | 0.897 (0.884) | 0.904 (0.897) | 0.901 (0.889) |
| (i)+(ii), reweighting by $\hat{w}$ (ProtCOVER-W) | 0.902 (0.898) | 0.903 (0.895) | 0.900 (0.893) | 0.896 (0.887) |
| (i)+(iii) gate | 0.879 (0.853) | 0.868 (0.820) | 0.904 (0.883) | 0.903 (0.897) |
| ProtCOVER-W + (iii) gate | 0.902 (0.898) | 0.906 (0.897) | 0.900 (0.894) | 0.896 (0.889) |
| (ii) without (i): variant-level weighting | 0.851 (0.821) | 0.852 (0.804) | 0.867 (0.819) | 0.862 (0.818) |
| *Descriptors for $\hat{w}$ (ProtCOVER-W)* | | | | |
| sequence similarity only | 0.876 (0.849) | 0.856 (0.805) | 0.899 (0.886) | 0.904 (0.900) |
| alignment depth only | 0.899 (0.890) | 0.899 (0.882) | 0.897 (0.879) | 0.902 (0.897) |
| embedding distance only | 0.882 (0.860) | 0.869 (0.822) | 0.904 (0.881) | 0.893 (0.883) |
| the three support groups | 0.902 (0.898) | 0.903 (0.895) | 0.900 (0.893) | 0.896 (0.888) |
| main set + predictor score summaries | 0.904 (0.895) | 0.902 (0.891) | 0.900 (0.887) | 0.894 (0.886) |
| *Weight clipping $c$ (default 20; ProtCOVER-W)* | | | | |
| $c = 5$ | 0.901 (0.896) | 0.891 (0.877) | 0.900 (0.893) | 0.904 (0.901) |
| $c = 100$ | 0.902 (0.898) | 0.909 (0.902) | 0.900 (0.893) | 0.889 (0.882) |
| no clipping | 0.902 (0.898) | 0.909 (0.902) | 0.900 (0.893) | 0.852 (0.824) |

**Table 5.** Robustness runs: worse-class shortfall below the 0.90 target at $\alpha = 0.1$, mean (maximum) over the 25 shifted cells of each run, class-conditional versions; columns as in Table 2.

| | | | ProtCOVER | | |
|---|---|---|---|---|---|
| Run | HCP | GWCP | -W | -D | **default** |
| Transductive base model | 0.023 (0.102) | 0.007 (0.027) | 0.003 (0.014) | 0.003 (0.015) | 0.001 (0.010) |
| Alignment-free descriptors | 0.019 (0.080) | 0.019 (0.080) | 0.015 (0.068) | 0.016 (0.070) | 0.011 (0.060) |
| Families: 50% identity | 0.021 (0.092) | 0.009 (0.041) | 0.003 (0.011) | 0.002 (0.016) | 0.001 (0.014) |
| Families: homology components | 0.017 (0.078) | 0.008 (0.028) | 0.003 (0.009) | 0.001 (0.007) | 0.001 (0.010) |
| Families: any significant hit | 0.023 (0.109) | 0.009 (0.037) | 0.002 (0.010) | 0.001 (0.008) | 0.001 (0.011) |

**Table 6.** Pooling, feature classes, method against information, and the share of calibration families that fit $\hat{w}$, over the 25 shifted cells at $\alpha = 0.1$ (class-conditional versions). Shortfall: worse-class shortfall, mean (maximum) over cells. Size: mean set size. s.d. $\text{Cov}_1$: seed-to-seed standard deviation of pathogenic coverage, mean over cells. Abst.: largest fraction of test families with both thresholds infinite in any cell. Margin: cells where ProtCOVER's shortfall is smaller / larger than the variant's by more than 0.01. Paired: cells where ProtCOVER's worse-class coverage is higher / lower than the variant's with a paired 95% interval that excludes zero. Variants were declared in the configuration before they were run.

| Variant | Shortfall | Size | s.d. $\text{Cov}_1$ | Abst. | Margin | Paired |
|---|---|---|---|---|---|---|
| ProtCOVER, features $(1, \hat{w})$, ratio fitted on a third (default) | 0.001 (0.008) | 1.294 | 0.016 | 0.3% | - | - |
| one variant per family and class (Gibbs et al.) | 0.002 (0.012) | 1.292 | 0.020 | 0.3% | 1/0 | 12/2 |
| features $(1, \hat{w}, d)$ | 0.000 (0.002) | 1.316 | 0.016 | 0.3% | 0/0 | 0/21 |
| ratio fitted on 20% of calibration families | 0.001 (0.011) | 1.286 | 0.017 | 1.0% | 0/0 | 11/2 |
| ratio fitted on 50% | 0.000 (0.003) | 1.315 | 0.018 | 1.5% | 0/0 | 2/12 |
| ProtCOVER-D, features $(1, d)$ (default) | 0.004 (0.016) | 1.252 | 0.016 | 0.0% | 5/0 | - |
| features $(1, d, d^2)$ | 0.001 (0.006) | 1.275 | 0.016 | 0.4% | 0/0 | 7/10 |
| ProtCOVER-W, ratio fitted on a third (default) | 0.003 (0.013) | 1.262 | 0.023 | 0.0% | 1/0 | 16/1 |
| ratio fitted on 20% | 0.003 (0.015) | 1.262 | 0.023 | 0.0% | 1/0 | - |
| ratio fitted on 50% | 0.003 (0.016) | 1.276 | 0.024 | 0.0% | 1/0 | - |
| GWCP, nine descriptor bins (baseline) | 0.007 (0.024) | 1.241 | 0.020 | 0.0% | 4/0 | 21/1 |
| bins at the ninths of $\hat{w}$ (ProtCOVER's information) | 0.006 (0.019) | 1.246 | 0.022 | 0.0% | 4/0 | 22/0 |
| Conditional calibration on GWCP's bins (GWCP's information) | 0.000 (0.000) | 1.312 | 0.017 | 13.4% | 0/0 | 1/12 |

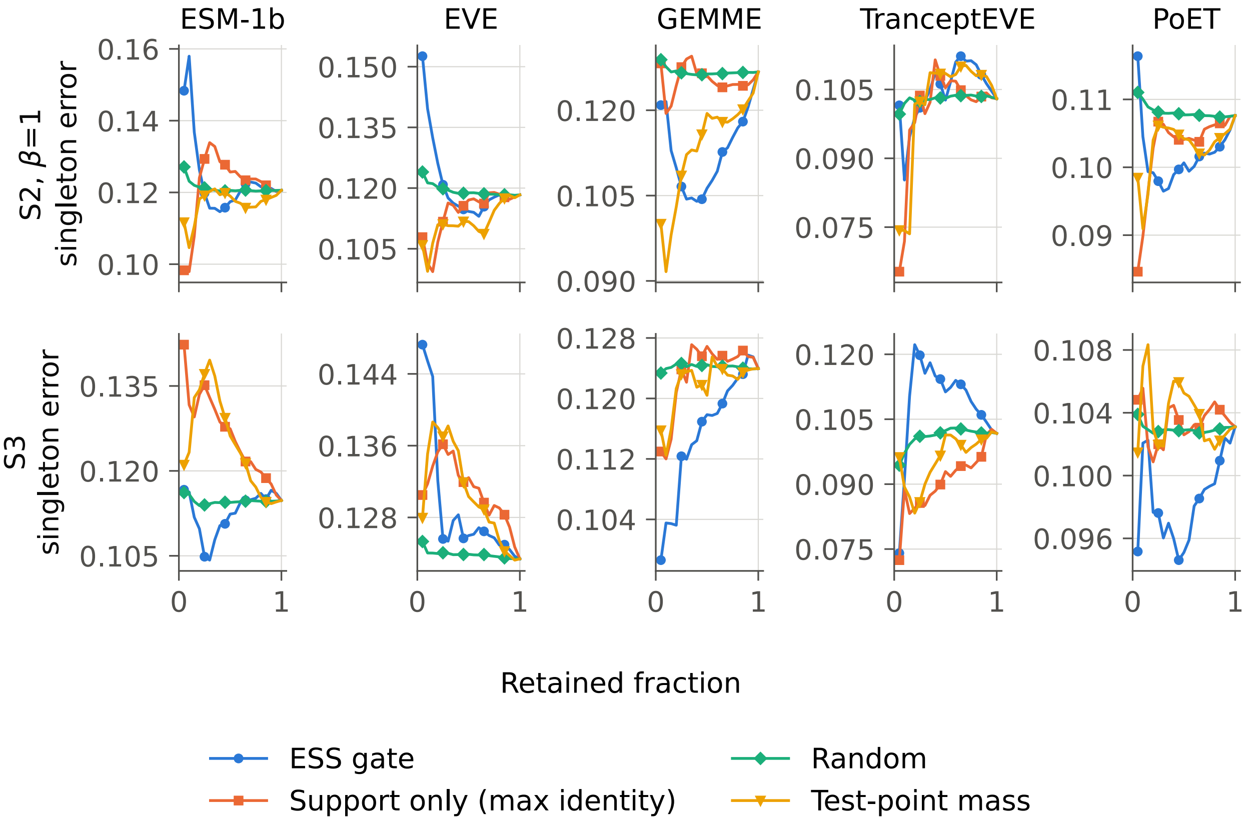


**Fig. 4.** Risk-coverage of abstention rules applied to ProtCOVER-W's class-conditional sets ($\alpha = 0.1$, mean over seeds): error of retained singleton sets as whole families are abstained in order of the score. Random abstention is averaged over 100 random family orders. The gate's decision depends only on label-free quantities, so it abstains on the same families whichever variant of (ii) it wraps.

**Table 7.** Paired differences in pathogenic coverage $\mathrm{Cov}_1$: ProtCOVER minus each baseline.

| Setting | Predictor | $\mathrm{Cov}_1$ (pathogenic) | | |
|---|---|---|---|---|
| | | − HCP | − GWCP | − ProtCOVER-W |
| S1 | ESM-1b | −0.001 [−0.004, +0.003] | +0.001 [−0.003, +0.004] | +0.004 [+0.000, +0.007] |
| | EVE | +0.001 [−0.002, +0.003] | +0.002 [−0.000, +0.004] | +0.000 [−0.002, +0.002] |
| | GEMME | +0.002 [+0.000, +0.003] | +0.002 [+0.001, +0.003] | +0.002 [+0.000, +0.003] |
| | TranceptEVE | +0.001 [−0.003, +0.006] | +0.001 [−0.003, +0.005] | +0.002 [−0.002, +0.007] |
| | PoET | +0.003 [+0.000, +0.005] | +0.004 [+0.001, +0.006] | +0.002 [+0.000, +0.005] |
| S2, $\beta = 0.5$ | ESM-1b | +0.021 [+0.017, +0.025] | −0.000 [−0.005, +0.004] | −0.001 [−0.005, +0.003] |
| | EVE | +0.013 [+0.010, +0.015] | +0.007 [+0.004, +0.009] | −0.001 [−0.003, +0.002] |
| | GEMME | +0.011 [+0.009, +0.013] | +0.010 [+0.007, +0.012] | +0.001 [−0.001, +0.003] |
| | TranceptEVE | +0.022 [+0.017, +0.026] | +0.006 [+0.002, +0.011] | −0.002 [−0.007, +0.002] |
| | PoET | +0.005 [+0.003, +0.007] | +0.011 [+0.009, +0.013] | +0.004 [+0.002, +0.006] |
| S2, $\beta = 1$ | ESM-1b | +0.057 [+0.051, +0.062] | +0.014 [+0.009, +0.019] | −0.004 [−0.009, +0.001] |
| | EVE | +0.022 [+0.019, +0.026] | +0.013 [+0.010, +0.016] | +0.004 [+0.001, +0.007] |
| | GEMME | +0.016 [+0.013, +0.019] | +0.015 [+0.013, +0.018] | +0.007 [+0.005, +0.010] |
| | TranceptEVE | +0.036 [+0.032, +0.041] | +0.004 [+0.000, +0.008] | +0.000 [−0.003, +0.004] |
| | PoET | −0.005 [−0.007, −0.004] | +0.006 [+0.004, +0.008] | +0.003 [+0.000, +0.004] |
| S2, $\beta = 2$ | ESM-1b | +0.104 [+0.098, +0.111] | +0.028 [+0.022, +0.034] | +0.018 [+0.013, +0.024] |
| | EVE | +0.049 [+0.045, +0.054] | +0.041 [+0.037, +0.046] | +0.025 [+0.021, +0.028] |
| | GEMME | +0.033 [+0.029, +0.037] | +0.032 [+0.028, +0.037] | +0.023 [+0.019, +0.027] |
| | TranceptEVE | +0.095 [+0.089, +0.102] | +0.060 [+0.054, +0.066] | +0.034 [+0.030, +0.039] |
| | PoET | +0.006 [+0.003, +0.009] | +0.015 [+0.012, +0.018] | +0.008 [+0.006, +0.011] |
| S3 | ESM-1b | +0.009 [+0.007, +0.011] | −0.002 [−0.004, −0.000] | −0.002 [−0.004, −0.000] |
| | EVE | −0.000 [−0.002, +0.001] | +0.002 [+0.000, +0.004] | +0.000 [−0.001, +0.001] |
| | GEMME | +0.004 [+0.002, +0.005] | +0.004 [+0.003, +0.006] | +0.000 [−0.001, +0.001] |
| | TranceptEVE | −0.010 [−0.012, −0.007] | +0.001 [−0.002, +0.004] | +0.000 [−0.002, +0.002] |
| | PoET | −0.017 [−0.020, −0.014] | +0.006 [+0.004, +0.009] | +0.006 [+0.004, +0.009] |
| S4 | ESM-1b | +0.016 [+0.013, +0.019] | +0.014 [+0.011, +0.018] | +0.025 [+0.022, +0.029] |
| | EVE | +0.009 [+0.007, +0.011] | +0.013 [+0.011, +0.016] | +0.008 [+0.005, +0.011] |
| | GEMME | +0.010 [+0.008, +0.012] | +0.013 [+0.010, +0.015] | +0.004 [+0.002, +0.006] |
| | TranceptEVE | +0.010 [+0.007, +0.013] | +0.015 [+0.011, +0.018] | +0.030 [+0.026, +0.035] |
| | PoET | +0.001 [−0.001, +0.003] | +0.008 [+0.005, +0.010] | +0.011 [+0.009, +0.014] |

**Table 8.** Paired differences in benign coverage $\mathrm{Cov}_0$: ProtCOVER minus each baseline.

| | | $\mathrm{Cov}_0$ (benign) | | |
|---|---|---|---|---|
| Setting | Predictor | − HCP | − GWCP | − ProtCOVER-W |
| S1 | ESM-1b | +0.005 [+0.003, +0.006] | +0.006 [+0.005, +0.008] | +0.001 [+0.000, +0.003] |
| | EVE | +0.006 [+0.004, +0.007] | +0.008 [+0.007, +0.010] | +0.003 [+0.002, +0.004] |
| | GEMME | +0.001 [−0.000, +0.002] | +0.003 [+0.001, +0.004] | +0.001 [−0.000, +0.002] |
| | TranceptEVE | +0.003 [−0.001, +0.006] | +0.004 [+0.001, +0.008] | +0.002 [−0.002, +0.005] |
| | PoET | +0.006 [+0.004, +0.008] | +0.008 [+0.006, +0.010] | +0.005 [+0.003, +0.006] |
| S2, $\beta = 0.5$ | ESM-1b | −0.004 [−0.005, −0.003] | +0.006 [+0.004, +0.007] | +0.003 [+0.002, +0.005] |
| | EVE | +0.002 [+0.001, +0.003] | +0.006 [+0.005, +0.007] | +0.002 [+0.001, +0.003] |
| | GEMME | +0.008 [+0.007, +0.009] | +0.009 [+0.008, +0.010] | +0.005 [+0.004, +0.005] |
| | TranceptEVE | −0.001 [−0.003, +0.001] | +0.000 [−0.002, +0.002] | +0.003 [+0.001, +0.005] |
| | PoET | +0.016 [+0.013, +0.018] | +0.006 [+0.004, +0.008] | −0.001 [−0.003, +0.001] |
| S2, $\beta = 1$ | ESM-1b | −0.011 [−0.012, −0.009] | +0.006 [+0.004, +0.008] | +0.004 [+0.003, +0.006] |
| | EVE | +0.001 [−0.000, +0.002] | +0.005 [+0.004, +0.006] | +0.003 [+0.002, +0.004] |
| | GEMME | +0.014 [+0.013, +0.015] | +0.016 [+0.014, +0.017] | +0.005 [+0.004, +0.006] |
| | TranceptEVE | +0.013 [+0.011, +0.015] | +0.014 [+0.012, +0.016] | +0.006 [+0.004, +0.008] |
| | PoET | +0.034 [+0.031, +0.038] | +0.017 [+0.014, +0.019] | +0.003 [+0.001, +0.006] |
| S2, $\beta = 2$ | ESM-1b | −0.007 [−0.008, −0.005] | +0.018 [+0.016, +0.020] | +0.008 [+0.006, +0.009] |
| | EVE | +0.017 [+0.015, +0.018] | +0.028 [+0.026, +0.030] | +0.012 [+0.010, +0.013] |
| | GEMME | +0.026 [+0.024, +0.028] | +0.044 [+0.041, +0.046] | +0.020 [+0.018, +0.022] |
| | TranceptEVE | +0.029 [+0.026, +0.033] | +0.032 [+0.029, +0.036] | +0.022 [+0.020, +0.026] |
| | PoET | +0.069 [+0.065, +0.073] | +0.048 [+0.044, +0.052] | +0.029 [+0.026, +0.032] |
| S3 | ESM-1b | +0.002 [+0.001, +0.003] | +0.007 [+0.006, +0.008] | +0.002 [+0.001, +0.002] |
| | EVE | +0.005 [+0.004, +0.006] | +0.003 [+0.003, +0.004] | +0.001 [+0.000, +0.001] |
| | GEMME | +0.003 [+0.002, +0.004] | +0.010 [+0.009, +0.011] | +0.001 [+0.001, +0.002] |
| | TranceptEVE | +0.028 [+0.026, +0.031] | +0.006 [+0.003, +0.008] | −0.004 [−0.006, −0.003] |
| | PoET | +0.020 [+0.018, +0.022] | +0.006 [+0.004, +0.008] | −0.004 [−0.005, −0.003] |
| S4 | ESM-1b | +0.003 [+0.002, +0.005] | +0.003 [+0.002, +0.005] | +0.001 [−0.001, +0.002] |
| | EVE | +0.003 [+0.002, +0.004] | +0.004 [+0.002, +0.005] | +0.005 [+0.004, +0.007] |
| | GEMME | +0.005 [+0.004, +0.006] | +0.008 [+0.006, +0.009] | +0.013 [+0.012, +0.015] |
| | TranceptEVE | +0.020 [+0.017, +0.023] | +0.012 [+0.009, +0.015] | −0.008 [−0.011, −0.005] |
| | PoET | +0.015 [+0.013, +0.017] | +0.011 [+0.009, +0.013] | +0.004 [+0.002, +0.006] |

**Table 9.** Paired differences in mean set size (variant-weighted): ProtCOVER minus each baseline.

| Setting | Predictor | Mean set size | | |
|---|---|---|---|---|
| | | − HCP | − GWCP | − ProtCOVER-W |
| S1 | ESM-1b | −0.002 [−0.006, +0.002] | +0.003 [−0.001, +0.007] | +0.004 [+0.000, +0.009] |
| | EVE | +0.014 [+0.012, +0.017] | +0.021 [+0.018, +0.023] | +0.007 [+0.005, +0.009] |
| | GEMME | +0.009 [+0.007, +0.011] | +0.012 [+0.009, +0.014] | +0.008 [+0.006, +0.010] |
| | TranceptEVE | −0.020 [−0.033, −0.008] | −0.018 [−0.030, −0.005] | −0.021 [−0.033, −0.008] |
| | PoET | +0.021 [+0.018, +0.025] | +0.026 [+0.022, +0.029] | +0.017 [+0.013, +0.020] |
| S2, $\beta = 0.5$ | ESM-1b | +0.033 [+0.025, +0.041] | −0.006 [−0.013, +0.001] | −0.011 [−0.018, −0.004] |
| | EVE | +0.020 [+0.017, +0.023] | +0.018 [+0.016, +0.021] | −0.002 [−0.004, +0.001] |
| | GEMME | +0.030 [+0.027, +0.033] | +0.030 [+0.028, +0.033] | +0.011 [+0.009, +0.014] |
| | TranceptEVE | +0.017 [+0.007, +0.028] | −0.010 [−0.021, +0.002] | −0.021 [−0.032, −0.009] |
| | PoET | +0.024 [+0.021, +0.027] | +0.022 [+0.020, +0.026] | −0.002 [−0.004, +0.001] |
| S2, $\beta = 1$ | ESM-1b | +0.092 [+0.082, +0.103] | +0.023 [+0.013, +0.032] | −0.018 [−0.027, −0.010] |
| | EVE | +0.037 [+0.032, +0.042] | +0.030 [+0.026, +0.035] | +0.012 [+0.008, +0.016] |
| | GEMME | +0.052 [+0.047, +0.058] | +0.053 [+0.048, +0.059] | +0.025 [+0.020, +0.030] |
| | TranceptEVE | +0.068 [+0.058, +0.080] | +0.017 [+0.005, +0.030] | +0.004 [−0.007, +0.018] |
| | PoET | +0.033 [+0.028, +0.039] | +0.032 [+0.027, +0.038] | +0.009 [+0.005, +0.014] |
| S2, $\beta = 2$ | ESM-1b | +0.207 [+0.192, +0.224] | +0.092 [+0.078, +0.106] | +0.059 [+0.049, +0.071] |
| | EVE | +0.169 [+0.156, +0.184] | +0.170 [+0.157, +0.185] | +0.114 [+0.104, +0.126] |
| | GEMME | +0.171 [+0.158, +0.187] | +0.184 [+0.170, +0.199] | +0.137 [+0.126, +0.150] |
| | TranceptEVE | +0.243 [+0.227, +0.261] | +0.191 [+0.171, +0.210] | +0.144 [+0.126, +0.164] |
| | PoET | +0.169 [+0.156, +0.184] | +0.159 [+0.145, +0.174] | +0.123 [+0.111, +0.135] |
| S3 | ESM-1b | +0.025 [+0.022, +0.028] | +0.007 [+0.004, +0.010] | −0.003 [−0.006, −0.000] |
| | EVE | +0.010 [+0.009, +0.012] | +0.012 [+0.011, +0.013] | +0.004 [+0.003, +0.005] |
| | GEMME | +0.014 [+0.012, +0.015] | +0.025 [+0.023, +0.026] | +0.003 [+0.002, +0.004] |
| | TranceptEVE | +0.033 [+0.024, +0.040] | +0.013 [+0.006, +0.019] | −0.006 [−0.011, −0.002] |
| | PoET | +0.002 [−0.000, +0.004] | +0.025 [+0.023, +0.026] | +0.008 [+0.007, +0.010] |
| S4 | ESM-1b | +0.046 [+0.036, +0.058] | +0.050 [+0.040, +0.063] | +0.064 [+0.054, +0.075] |
| | EVE | +0.036 [+0.029, +0.045] | +0.043 [+0.036, +0.052] | +0.026 [+0.020, +0.032] |
| | GEMME | +0.040 [+0.034, +0.046] | +0.047 [+0.041, +0.053] | +0.037 [+0.031, +0.042] |
| | TranceptEVE | +0.041 [+0.031, +0.051] | +0.043 [+0.034, +0.053] | +0.038 [+0.024, +0.051] |
| | PoET | +0.036 [+0.031, +0.043] | +0.046 [+0.040, +0.052] | +0.035 [+0.030, +0.040] |

**Table 10.** Paired differences in singleton error: ProtCOVER minus each baseline.

| | | Singleton error | | |
|---|---|---|---|---|
| Setting | Predictor | − HCP | − GWCP | − ProtCOVER-W |
| S1 | ESM-1b | −0.000 [−0.002, +0.001] | −0.001 [−0.003, +0.000] | +0.001 [−0.001, +0.002] |
| | EVE | −0.001 [−0.002, +0.000] | −0.003 [−0.004, −0.002] | +0.000 [−0.001, +0.002] |
| | GEMME | −0.004 [−0.004, −0.003] | −0.004 [−0.005, −0.003] | −0.003 [−0.004, −0.002] |
| | TranceptEVE | +0.011 [+0.003, +0.019] | +0.011 [+0.003, +0.019] | +0.010 [+0.002, +0.018] |
| | PoET | −0.002 [−0.004, −0.000] | −0.003 [−0.005, −0.001] | −0.001 [−0.002, +0.001] |
| S2, $\beta = 0.5$ | ESM-1b | +0.005 [+0.003, +0.008] | +0.000 [−0.002, +0.003] | +0.001 [−0.002, +0.003] |
| | EVE | −0.000 [−0.001, +0.001] | −0.001 [−0.002, +0.000] | +0.004 [+0.003, +0.005] |
| | GEMME | −0.006 [−0.006, −0.005] | −0.006 [−0.007, −0.005] | −0.002 [−0.003, −0.001] |
| | TranceptEVE | +0.015 [+0.006, +0.024] | +0.028 [+0.017, +0.038] | +0.032 [+0.020, +0.043] |
| | PoET | −0.009 [−0.010, −0.007] | −0.006 [−0.007, −0.004] | −0.001 [−0.002, +0.001] |
| S2, $\beta = 1$ | ESM-1b | +0.006 [+0.004, +0.010] | −0.001 [−0.004, +0.002] | −0.000 [−0.004, +0.003] |
| | EVE | −0.002 [−0.003, −0.001] | −0.002 [−0.003, −0.000] | +0.002 [+0.001, +0.003] |
| | GEMME | −0.010 [−0.011, −0.008] | −0.010 [−0.012, −0.009] | −0.003 [−0.005, −0.002] |
| | TranceptEVE | +0.005 [−0.003, +0.012] | +0.028 [+0.016, +0.039] | +0.028 [+0.017, +0.039] |
| | PoET | −0.016 [−0.018, −0.014] | −0.010 [−0.011, −0.008] | −0.003 [−0.005, −0.001] |
| S2, $\beta = 2$ | ESM-1b | +0.003 [−0.000, +0.007] | −0.010 [−0.013, −0.006] | −0.002 [−0.005, +0.002] |
| | EVE | −0.007 [−0.009, −0.004] | −0.011 [−0.014, −0.009] | +0.001 [−0.001, +0.003] |
| | GEMME | −0.010 [−0.013, −0.007] | −0.021 [−0.024, −0.018] | −0.005 [−0.008, −0.003] |
| | TranceptEVE | −0.005 [−0.015, +0.006] | +0.020 [+0.004, +0.035] | +0.032 [+0.018, +0.046] |
| | PoET | −0.027 [−0.030, −0.024] | −0.019 [−0.022, −0.016] | −0.011 [−0.013, −0.008] |
| S3 | ESM-1b | −0.003 [−0.003, −0.002] | −0.005 [−0.006, −0.004] | −0.002 [−0.002, −0.001] |
| | EVE | −0.002 [−0.003, −0.001] | −0.002 [−0.003, −0.002] | −0.000 [−0.001, +0.000] |
| | GEMME | −0.003 [−0.003, −0.002] | −0.006 [−0.007, −0.005] | −0.001 [−0.001, −0.000] |
| | TranceptEVE | +0.005 [−0.002, +0.013] | −0.000 [−0.007, +0.006] | +0.002 [−0.002, +0.005] |
| | PoET | −0.002 [−0.004, −0.001] | −0.003 [−0.004, −0.001] | +0.002 [+0.001, +0.003] |
| S4 | ESM-1b | +0.001 [−0.002, +0.003] | −0.001 [−0.003, +0.001] | −0.003 [−0.005, −0.001] |
| | EVE | −0.001 [−0.002, +0.001] | −0.004 [−0.006, −0.003] | −0.005 [−0.007, −0.003] |
| | GEMME | −0.004 [−0.005, −0.003] | −0.006 [−0.008, −0.005] | −0.006 [−0.007, −0.004] |
| | TranceptEVE | +0.005 [−0.004, +0.014] | −0.007 [−0.015, +0.001] | −0.032 [−0.045, −0.019] |
| | PoET | −0.004 [−0.006, −0.002] | −0.005 [−0.006, −0.003] | −0.004 [−0.006, −0.003] |

Tables 11–13 report every method, marginal (cc = no) and class-conditional (cc = yes), averaged over the five predictors: family-weighted coverage Cov, variant-weighted coverage, $Cov_1$, $Cov_0$, mean set size, singleton rate, the fraction of families with at least ten variants whose coverage is below $1 - \alpha - 0.05$, and the fraction of test families on which the method abstained. Per-predictor values with bootstrap intervals, and the summaries of every sensitivity, stress and external run, are in `results/` of the code repository.

Table 11: All methods at $\alpha = 0.1$, mean over predictors and seeds.

| Method | cc | Cov | var.-w. | $Cov_1$ | $Cov_0$ | size | single. | under. | abst. |
|---|---|---|---|---|---|---|---|---|---|
| *S1 (exchangeable)* | | | | | | | | | |
| Split (B1/B2) | no | 0.892 | 0.896 | 0.863 | 0.893 | 1.174 | 0.826 | 0.236 | 0.000 |
| Split (B1/B2) | yes | 0.890 | 0.896 | 0.861 | 0.891 | 1.172 | 0.828 | 0.241 | 0.000 |
| HCP (B3) | no | 0.900 | 0.908 | 0.874 | 0.902 | 1.207 | 0.793 | 0.210 | 0.000 |
| HCP (B3) | yes | 0.909 | 0.914 | 0.902 | 0.900 | 1.241 | 0.759 | 0.197 | 0.000 |
| Subsample once (B3) | no | 0.899 | 0.907 | 0.874 | 0.901 | 1.204 | 0.796 | 0.215 | 0.000 |
| Subsample once (B3) | yes | 0.908 | 0.914 | 0.903 | 0.898 | 1.242 | 0.758 | 0.199 | 0.000 |
| Repeated subs. (B3) | no | 0.900 | 0.908 | 0.874 | 0.902 | 1.207 | 0.793 | 0.210 | 0.000 |
| Repeated subs. (B3) | yes | 0.908 | 0.913 | 0.902 | 0.899 | 1.240 | 0.760 | 0.199 | 0.000 |
| GWCP (B4) | no | 0.899 | 0.906 | 0.872 | 0.901 | 1.201 | 0.799 | 0.216 | 0.000 |
| GWCP (B4) | yes | 0.907 | 0.913 | 0.902 | 0.898 | 1.237 | 0.763 | 0.202 | 0.000 |
| ProtCOVER | no | 0.903 | 0.906 | 0.872 | 0.907 | 1.206 | 0.794 | 0.204 | 0.000 |
| ProtCOVER | yes | 0.910 | 0.914 | 0.903 | 0.904 | 1.246 | 0.754 | 0.196 | 0.000 |
| ProtCOVER-D | no | 0.906 | 0.897 | 0.868 | 0.914 | 1.196 | 0.804 | 0.202 | 0.000 |
| ProtCOVER-D | yes | 0.909 | 0.912 | 0.908 | 0.905 | 1.242 | 0.758 | 0.207 | 0.000 |
| ProtCOVER-W | no | 0.901 | 0.908 | 0.874 | 0.902 | 1.208 | 0.792 | 0.209 | 0.000 |
| ProtCOVER-W | yes | 0.909 | 0.914 | 0.901 | 0.902 | 1.243 | 0.757 | 0.194 | 0.000 |
| (i)+(iii) gate | no | 0.900 | 0.908 | 0.874 | 0.902 | 1.207 | 0.793 | 0.209 | 0.000 |
| (i)+(iii) gate | yes | 0.909 | 0.914 | 0.902 | 0.900 | 1.242 | 0.758 | 0.197 | 0.001 |
| ProtCOVER-W + gate | no | 0.901 | 0.908 | 0.874 | 0.903 | 1.208 | 0.792 | 0.209 | 0.001 |
| ProtCOVER-W + gate | yes | 0.910 | 0.914 | 0.901 | 0.902 | 1.243 | 0.757 | 0.194 | 0.000 |
| Support-only (B5) | no | 0.901 | 0.908 | 0.874 | 0.902 | 1.208 | 0.792 | 0.209 | 0.001 |
| Support-only (B5) | yes | 0.909 | 0.914 | 0.901 | 0.902 | 1.243 | 0.757 | 0.194 | 0.000 |
| Random abst. (B5) | no | 0.901 | 0.908 | 0.874 | 0.902 | 1.208 | 0.792 | 0.209 | 0.001 |
| Random abst. (B5) | yes | 0.909 | 0.914 | 0.901 | 0.902 | 1.243 | 0.757 | 0.194 | 0.000 |
| *S2*, $\beta = 0.5$ | | | | | | | | | |
| Split (B1/B2) | no | 0.889 | 0.892 | 0.854 | 0.892 | 1.170 | 0.830 | 0.248 | 0.000 |
| Split (B1/B2) | yes | 0.888 | 0.891 | 0.848 | 0.893 | 1.168 | 0.832 | 0.252 | 0.000 |
| HCP (B3) | no | 0.897 | 0.904 | 0.866 | 0.900 | 1.202 | 0.798 | 0.223 | 0.000 |
| HCP (B3) | yes | 0.904 | 0.908 | 0.890 | 0.898 | 1.229 | 0.771 | 0.215 | 0.000 |
| Subsample once (B3) | no | 0.896 | 0.902 | 0.866 | 0.897 | 1.198 | 0.802 | 0.232 | 0.000 |
| Subsample once (B3) | yes | 0.904 | 0.908 | 0.892 | 0.897 | 1.231 | 0.769 | 0.216 | 0.000 |
| Repeated subs. (B3) | no | 0.897 | 0.904 | 0.866 | 0.900 | 1.201 | 0.799 | 0.224 | 0.000 |
| Repeated subs. (B3) | yes | 0.904 | 0.908 | 0.890 | 0.898 | 1.229 | 0.771 | 0.215 | 0.000 |
| GWCP (B4) | no | 0.898 | 0.904 | 0.866 | 0.901 | 1.203 | 0.797 | 0.222 | 0.000 |
| GWCP (B4) | yes | 0.906 | 0.911 | 0.898 | 0.897 | 1.242 | 0.758 | 0.211 | 0.000 |
| ProtCOVER | no | 0.902 | 0.908 | 0.872 | 0.905 | 1.217 | 0.783 | 0.208 | 0.000 |
| ProtCOVER | yes | 0.910 | 0.911 | 0.905 | 0.903 | 1.253 | 0.747 | 0.198 | 0.000 |
| ProtCOVER-D | no | 0.904 | 0.896 | 0.861 | 0.913 | 1.198 | 0.802 | 0.209 | 0.000 |
| ProtCOVER-D | yes | 0.908 | 0.911 | 0.905 | 0.905 | 1.250 | 0.750 | 0.213 | 0.000 |
| ProtCOVER-W | no | 0.901 | 0.907 | 0.870 | 0.904 | 1.214 | 0.786 | 0.211 | 0.000 |
| ProtCOVER-W | yes | 0.910 | 0.915 | 0.904 | 0.900 | 1.258 | 0.742 | 0.196 | 0.000 |
| (i)+(iii) gate | no | 0.897 | 0.904 | 0.866 | 0.900 | 1.202 | 0.798 | 0.223 | 0.000 |

*All methods at $\alpha = 0.1$, continued*

| Method | cc | Cov | var.-w. | $\mathrm{Cov}_1$ | $\mathrm{Cov}_0$ | size | single. | under. | abst. |
|---|---|---|---|---|---|---|---|---|---|
| (i)+(iii) gate | yes | 0.904 | 0.908 | 0.890 | 0.899 | 1.229 | 0.771 | 0.215 | 0.001 |
| ProtCOVER-W + gate | no | 0.901 | 0.907 | 0.870 | 0.904 | 1.215 | 0.785 | 0.211 | 0.001 |
| ProtCOVER-W + gate | yes | 0.910 | 0.915 | 0.904 | 0.900 | 1.259 | 0.741 | 0.196 | 0.000 |
| Support-only (B5) | no | 0.901 | 0.907 | 0.870 | 0.904 | 1.214 | 0.786 | 0.211 | 0.001 |
| Support-only (B5) | yes | 0.910 | 0.915 | 0.904 | 0.900 | 1.259 | 0.741 | 0.196 | 0.000 |
| Random abst. (B5) | no | 0.901 | 0.907 | 0.870 | 0.904 | 1.215 | 0.785 | 0.211 | 0.001 |
| Random abst. (B5) | yes | 0.910 | 0.915 | 0.904 | 0.900 | 1.259 | 0.741 | 0.196 | 0.000 |
| *S2, $\beta = 1$* | | | | | | | | | |
| Split (B1/B2) | no | 0.887 | 0.890 | 0.847 | 0.891 | 1.168 | 0.832 | 0.257 | 0.000 |
| Split (B1/B2) | yes | 0.886 | 0.889 | 0.838 | 0.894 | 1.166 | 0.834 | 0.261 | 0.000 |
| HCP (B3) | no | 0.895 | 0.902 | 0.860 | 0.900 | 1.199 | 0.801 | 0.231 | 0.000 |
| HCP (B3) | yes | 0.901 | 0.904 | 0.879 | 0.898 | 1.219 | 0.781 | 0.226 | 0.000 |
| Subsample once (B3) | no | 0.895 | 0.902 | 0.859 | 0.900 | 1.200 | 0.800 | 0.232 | 0.000 |
| Subsample once (B3) | yes | 0.902 | 0.904 | 0.879 | 0.899 | 1.223 | 0.777 | 0.223 | 0.000 |
| Repeated subs. (B3) | no | 0.896 | 0.902 | 0.859 | 0.900 | 1.199 | 0.801 | 0.231 | 0.000 |
| Repeated subs. (B3) | yes | 0.901 | 0.904 | 0.879 | 0.898 | 1.219 | 0.781 | 0.227 | 0.000 |
| GWCP (B4) | no | 0.897 | 0.903 | 0.861 | 0.901 | 1.204 | 0.796 | 0.227 | 0.000 |
| GWCP (B4) | yes | 0.905 | 0.909 | 0.893 | 0.897 | 1.245 | 0.755 | 0.216 | 0.000 |
| ProtCOVER | no | 0.908 | 0.913 | 0.880 | 0.910 | 1.248 | 0.752 | 0.198 | 0.000 |
| ProtCOVER | yes | 0.914 | 0.914 | 0.904 | 0.909 | 1.276 | 0.724 | 0.193 | 0.000 |
| ProtCOVER-D | no | 0.903 | 0.896 | 0.855 | 0.914 | 1.201 | 0.799 | 0.214 | 0.000 |
| ProtCOVER-D | yes | 0.909 | 0.911 | 0.899 | 0.907 | 1.255 | 0.745 | 0.211 | 0.000 |
| ProtCOVER-W | no | 0.903 | 0.910 | 0.870 | 0.906 | 1.225 | 0.775 | 0.206 | 0.000 |
| ProtCOVER-W | yes | 0.912 | 0.915 | 0.902 | 0.905 | 1.269 | 0.731 | 0.195 | 0.000 |
| (i)+(iii) gate | no | 0.896 | 0.902 | 0.860 | 0.900 | 1.199 | 0.801 | 0.231 | 0.000 |
| (i)+(iii) gate | yes | 0.901 | 0.904 | 0.879 | 0.899 | 1.220 | 0.780 | 0.226 | 0.001 |
| ProtCOVER-W + gate | no | 0.903 | 0.910 | 0.871 | 0.907 | 1.225 | 0.775 | 0.206 | 0.001 |
| ProtCOVER-W + gate | yes | 0.912 | 0.915 | 0.902 | 0.905 | 1.269 | 0.731 | 0.195 | 0.000 |
| Support-only (B5) | no | 0.903 | 0.910 | 0.871 | 0.907 | 1.225 | 0.775 | 0.206 | 0.001 |
| Support-only (B5) | yes | 0.912 | 0.915 | 0.902 | 0.905 | 1.269 | 0.731 | 0.195 | 0.000 |
| Random abst. (B5) | no | 0.903 | 0.910 | 0.871 | 0.907 | 1.225 | 0.775 | 0.206 | 0.001 |
| Random abst. (B5) | yes | 0.912 | 0.915 | 0.902 | 0.905 | 1.269 | 0.731 | 0.195 | 0.000 |
| *S2, $\beta = 2$* | | | | | | | | | |
| Split (B1/B2) | no | 0.886 | 0.889 | 0.842 | 0.890 | 1.167 | 0.833 | 0.263 | 0.000 |
| Split (B1/B2) | yes | 0.884 | 0.888 | 0.833 | 0.893 | 1.165 | 0.835 | 0.269 | 0.000 |
| HCP (B3) | no | 0.894 | 0.900 | 0.854 | 0.898 | 1.196 | 0.804 | 0.239 | 0.000 |
| HCP (B3) | yes | 0.896 | 0.898 | 0.868 | 0.895 | 1.208 | 0.792 | 0.243 | 0.000 |
| Subsample once (B3) | no | 0.890 | 0.897 | 0.854 | 0.893 | 1.190 | 0.810 | 0.251 | 0.000 |
| Subsample once (B3) | yes | 0.894 | 0.896 | 0.866 | 0.892 | 1.203 | 0.797 | 0.251 | 0.000 |
| Repeated subs. (B3) | no | 0.893 | 0.900 | 0.854 | 0.898 | 1.194 | 0.806 | 0.241 | 0.000 |
| Repeated subs. (B3) | yes | 0.896 | 0.898 | 0.868 | 0.895 | 1.208 | 0.792 | 0.243 | 0.000 |
| GWCP (B4) | no | 0.894 | 0.900 | 0.855 | 0.899 | 1.201 | 0.799 | 0.242 | 0.000 |
| GWCP (B4) | yes | 0.898 | 0.902 | 0.890 | 0.888 | 1.241 | 0.759 | 0.243 | 0.000 |
| ProtCOVER | no | 0.926 | 0.931 | 0.908 | 0.927 | 1.387 | 0.613 | 0.156 | 0.000 |
| ProtCOVER | yes | 0.928 | 0.929 | 0.925 | 0.922 | 1.400 | 0.600 | 0.160 | 0.000 |
| ProtCOVER-D | no | 0.900 | 0.896 | 0.850 | 0.912 | 1.203 | 0.797 | 0.221 | 0.000 |
| ProtCOVER-D | yes | 0.907 | 0.909 | 0.897 | 0.905 | 1.270 | 0.730 | 0.220 | 0.000 |
| ProtCOVER-W | no | 0.909 | 0.915 | 0.880 | 0.912 | 1.259 | 0.741 | 0.192 | 0.000 |
| ProtCOVER-W | yes | 0.912 | 0.916 | 0.903 | 0.904 | 1.284 | 0.716 | 0.195 | 0.000 |

*All methods at $\alpha = 0.1$, continued*

| Method | cc | Cov | var.-w. | $Cov_1$ | $Cov_0$ | size | single. | under. | abst. |
|---|---|---|---|---|---|---|---|---|---|
| (i)+(iii) gate | no | 0.894 | 0.900 | 0.854 | 0.898 | 1.196 | 0.804 | 0.239 | 0.000 |
| (i)+(iii) gate | yes | 0.896 | 0.899 | 0.868 | 0.895 | 1.209 | 0.791 | 0.242 | 0.001 |
| ProtCOVER-W + gate | no | 0.910 | 0.916 | 0.881 | 0.912 | 1.264 | 0.736 | 0.190 | 0.007 |
| ProtCOVER-W + gate | yes | 0.914 | 0.918 | 0.906 | 0.906 | 1.301 | 0.699 | 0.191 | 0.028 |
| Support-only (B5) | no | 0.909 | 0.916 | 0.881 | 0.912 | 1.263 | 0.737 | 0.190 | 0.007 |
| Support-only (B5) | yes | 0.914 | 0.917 | 0.905 | 0.907 | 1.300 | 0.700 | 0.192 | 0.028 |
| Random abst. (B5) | no | 0.910 | 0.916 | 0.881 | 0.912 | 1.264 | 0.736 | 0.190 | 0.007 |
| Random abst. (B5) | yes | 0.914 | 0.918 | 0.905 | 0.907 | 1.302 | 0.698 | 0.191 | 0.028 |
| *S3 (orphan families)* | | | | | | | | | |
| Split (B1/B2) | no | 0.890 | 0.893 | 0.863 | 0.890 | 1.174 | 0.826 | 0.248 | 0.000 |
| Split (B1/B2) | yes | 0.888 | 0.893 | 0.860 | 0.889 | 1.171 | 0.829 | 0.251 | 0.000 |
| HCP (B3) | no | 0.899 | 0.904 | 0.875 | 0.900 | 1.209 | 0.791 | 0.219 | 0.000 |
| HCP (B3) | yes | 0.910 | 0.913 | 0.904 | 0.900 | 1.244 | 0.756 | 0.198 | 0.000 |
| Subsample once (B3) | no | 0.898 | 0.903 | 0.873 | 0.900 | 1.207 | 0.793 | 0.225 | 0.000 |
| Subsample once (B3) | yes | 0.909 | 0.912 | 0.903 | 0.899 | 1.243 | 0.757 | 0.202 | 0.000 |
| Repeated subs. (B3) | no | 0.899 | 0.904 | 0.875 | 0.900 | 1.208 | 0.792 | 0.221 | 0.000 |
| Repeated subs. (B3) | yes | 0.910 | 0.913 | 0.904 | 0.900 | 1.244 | 0.756 | 0.199 | 0.000 |
| GWCP (B4) | no | 0.902 | 0.908 | 0.880 | 0.904 | 1.226 | 0.774 | 0.210 | 0.000 |
| GWCP (B4) | yes | 0.911 | 0.911 | 0.899 | 0.905 | 1.245 | 0.755 | 0.202 | 0.000 |
| ProtCOVER | no | 0.907 | 0.912 | 0.886 | 0.908 | 1.244 | 0.756 | 0.196 | 0.000 |
| ProtCOVER | yes | 0.915 | 0.916 | 0.901 | 0.912 | 1.261 | 0.739 | 0.185 | 0.000 |
| ProtCOVER-D | no | 0.909 | 0.904 | 0.874 | 0.917 | 1.219 | 0.781 | 0.201 | 0.000 |
| ProtCOVER-D | yes | 0.912 | 0.910 | 0.904 | 0.911 | 1.244 | 0.756 | 0.208 | 0.000 |
| ProtCOVER-W | no | 0.907 | 0.912 | 0.886 | 0.909 | 1.244 | 0.756 | 0.197 | 0.000 |
| ProtCOVER-W | yes | 0.916 | 0.916 | 0.900 | 0.913 | 1.260 | 0.740 | 0.185 | 0.000 |
| (i)+(iii) gate | no | 0.899 | 0.904 | 0.875 | 0.900 | 1.209 | 0.791 | 0.219 | 0.000 |
| (i)+(iii) gate | yes | 0.910 | 0.913 | 0.904 | 0.901 | 1.245 | 0.755 | 0.198 | 0.002 |
| ProtCOVER-W + gate | no | 0.907 | 0.912 | 0.886 | 0.909 | 1.244 | 0.756 | 0.197 | 0.000 |
| ProtCOVER-W + gate | yes | 0.916 | 0.916 | 0.900 | 0.913 | 1.261 | 0.739 | 0.185 | 0.002 |
| Support-only (B5) | no | 0.907 | 0.912 | 0.886 | 0.909 | 1.244 | 0.756 | 0.197 | 0.000 |
| Support-only (B5) | yes | 0.916 | 0.916 | 0.900 | 0.913 | 1.261 | 0.739 | 0.185 | 0.002 |
| Random abst. (B5) | no | 0.907 | 0.912 | 0.886 | 0.909 | 1.244 | 0.756 | 0.197 | 0.000 |
| Random abst. (B5) | yes | 0.916 | 0.916 | 0.900 | 0.913 | 1.261 | 0.739 | 0.185 | 0.002 |
| *S4 (held-out regions)* | | | | | | | | | |
| Split (B1/B2) | no | 0.891 | 0.895 | 0.864 | 0.895 | 1.182 | 0.818 | 0.236 | 0.000 |
| Split (B1/B2) | yes | 0.889 | 0.896 | 0.864 | 0.890 | 1.177 | 0.823 | 0.242 | 0.000 |
| HCP (B3) | no | 0.898 | 0.904 | 0.874 | 0.902 | 1.208 | 0.792 | 0.216 | 0.000 |
| HCP (B3) | yes | 0.908 | 0.914 | 0.902 | 0.899 | 1.240 | 0.760 | 0.198 | 0.000 |
| Subsample once (B3) | no | 0.898 | 0.903 | 0.875 | 0.901 | 1.206 | 0.794 | 0.218 | 0.000 |
| Subsample once (B3) | yes | 0.907 | 0.913 | 0.901 | 0.899 | 1.239 | 0.761 | 0.199 | 0.000 |
| Repeated subs. (B3) | no | 0.898 | 0.903 | 0.873 | 0.902 | 1.208 | 0.792 | 0.216 | 0.000 |
| Repeated subs. (B3) | yes | 0.908 | 0.914 | 0.903 | 0.899 | 1.240 | 0.760 | 0.197 | 0.000 |
| GWCP (B4) | no | 0.898 | 0.904 | 0.875 | 0.903 | 1.210 | 0.790 | 0.216 | 0.000 |
| GWCP (B4) | yes | 0.907 | 0.910 | 0.899 | 0.901 | 1.234 | 0.766 | 0.204 | 0.000 |
| ProtCOVER | no | 0.907 | 0.911 | 0.883 | 0.912 | 1.250 | 0.750 | 0.197 | 0.003 |
| ProtCOVER | yes | 0.915 | 0.919 | 0.912 | 0.908 | 1.280 | 0.720 | 0.184 | 0.003 |
| ProtCOVER-D | no | 0.907 | 0.898 | 0.872 | 0.918 | 1.213 | 0.787 | 0.201 | 0.000 |
| ProtCOVER-D | yes | 0.908 | 0.909 | 0.901 | 0.909 | 1.241 | 0.759 | 0.212 | 0.000 |
| ProtCOVER-W | no | 0.905 | 0.910 | 0.881 | 0.910 | 1.240 | 0.760 | 0.199 | 0.000 |

*All methods at $\alpha = 0.1$, continued*

| Method | cc | Cov | var.-w. | $Cov_1$ | $Cov_0$ | size | single. | under. | abst. |
|---|---|---|---|---|---|---|---|---|---|
| ProtCOVER-W | yes | 0.908 | 0.906 | 0.896 | 0.905 | 1.240 | 0.760 | 0.210 | 0.000 |
| (i)+(iii) gate | no | 0.898 | 0.904 | 0.874 | 0.902 | 1.208 | 0.792 | 0.215 | 0.001 |
| (i)+(iii) gate | yes | 0.908 | 0.914 | 0.903 | 0.899 | 1.242 | 0.758 | 0.197 | 0.001 |
| ProtCOVER-W + gate | no | 0.905 | 0.910 | 0.882 | 0.910 | 1.240 | 0.760 | 0.199 | 0.001 |
| ProtCOVER-W + gate | yes | 0.908 | 0.907 | 0.896 | 0.905 | 1.243 | 0.757 | 0.209 | 0.005 |
| Support-only (B5) | no | 0.905 | 0.910 | 0.882 | 0.910 | 1.240 | 0.760 | 0.198 | 0.001 |
| Support-only (B5) | yes | 0.908 | 0.907 | 0.896 | 0.906 | 1.243 | 0.757 | 0.209 | 0.005 |
| Random abst. (B5) | no | 0.905 | 0.910 | 0.882 | 0.910 | 1.240 | 0.760 | 0.199 | 0.001 |
| Random abst. (B5) | yes | 0.908 | 0.907 | 0.896 | 0.906 | 1.243 | 0.757 | 0.209 | 0.005 |

Table 12: All methods at $\alpha = 0.05$, mean over predictors and seeds.

| Method | cc | Cov | var.-w. | $Cov_1$ | $Cov_0$ | size | single. | under. | abst. |
|---|---|---|---|---|---|---|---|---|---|
| *S1 (exchangeable)* | | | | | | | | | |
| Split (B1/B2) | no | 0.941 | 0.949 | 0.925 | 0.939 | 1.372 | 0.628 | 0.149 | 0.000 |
| Split (B1/B2) | yes | 0.943 | 0.948 | 0.920 | 0.945 | 1.371 | 0.629 | 0.145 | 0.000 |
| HCP (B3) | no | 0.951 | 0.958 | 0.938 | 0.951 | 1.445 | 0.555 | 0.117 | 0.000 |
| HCP (B3) | yes | 0.956 | 0.959 | 0.952 | 0.951 | 1.454 | 0.546 | 0.115 | 0.000 |
| Subsample once (B3) | no | 0.950 | 0.956 | 0.937 | 0.950 | 1.438 | 0.562 | 0.124 | 0.000 |
| Subsample once (B3) | yes | 0.956 | 0.959 | 0.953 | 0.951 | 1.455 | 0.545 | 0.116 | 0.000 |
| Repeated subs. (B3) | no | 0.951 | 0.958 | 0.938 | 0.951 | 1.446 | 0.554 | 0.117 | 0.000 |
| Repeated subs. (B3) | yes | 0.956 | 0.959 | 0.952 | 0.951 | 1.453 | 0.547 | 0.116 | 0.000 |
| GWCP (B4) | no | 0.949 | 0.956 | 0.936 | 0.950 | 1.434 | 0.566 | 0.124 | 0.000 |
| GWCP (B4) | yes | 0.955 | 0.958 | 0.951 | 0.949 | 1.446 | 0.554 | 0.121 | 0.000 |
| ProtCOVER | no | 0.954 | 0.958 | 0.938 | 0.956 | 1.440 | 0.560 | 0.112 | 0.000 |
| ProtCOVER | yes | 0.959 | 0.961 | 0.955 | 0.954 | 1.467 | 0.533 | 0.109 | 0.000 |
| ProtCOVER-D | no | 0.956 | 0.953 | 0.933 | 0.961 | 1.424 | 0.576 | 0.110 | 0.000 |
| ProtCOVER-D | yes | 0.958 | 0.961 | 0.957 | 0.956 | 1.478 | 0.522 | 0.112 | 0.000 |
| ProtCOVER-W | no | 0.952 | 0.958 | 0.939 | 0.952 | 1.449 | 0.551 | 0.115 | 0.000 |
| ProtCOVER-W | yes | 0.957 | 0.960 | 0.953 | 0.952 | 1.461 | 0.539 | 0.111 | 0.000 |
| (i)+(iii) gate | no | 0.951 | 0.958 | 0.938 | 0.951 | 1.445 | 0.555 | 0.117 | 0.000 |
| (i)+(iii) gate | yes | 0.956 | 0.959 | 0.952 | 0.951 | 1.454 | 0.546 | 0.115 | 0.000 |
| ProtCOVER-W + gate | no | 0.952 | 0.958 | 0.939 | 0.952 | 1.449 | 0.551 | 0.115 | 0.000 |
| ProtCOVER-W + gate | yes | 0.957 | 0.960 | 0.953 | 0.952 | 1.461 | 0.539 | 0.111 | 0.000 |
| Support-only (B5) | no | 0.952 | 0.958 | 0.939 | 0.952 | 1.449 | 0.551 | 0.115 | 0.000 |
| Support-only (B5) | yes | 0.957 | 0.960 | 0.953 | 0.952 | 1.461 | 0.539 | 0.111 | 0.000 |
| Random abst. (B5) | no | 0.952 | 0.958 | 0.939 | 0.952 | 1.449 | 0.551 | 0.115 | 0.000 |
| Random abst. (B5) | yes | 0.957 | 0.960 | 0.953 | 0.952 | 1.461 | 0.539 | 0.111 | 0.000 |
| *S2, $\beta = 0.5$* | | | | | | | | | |
| Split (B1/B2) | no | 0.938 | 0.946 | 0.918 | 0.937 | 1.366 | 0.634 | 0.162 | 0.000 |
| Split (B1/B2) | yes | 0.940 | 0.945 | 0.910 | 0.945 | 1.364 | 0.636 | 0.158 | 0.000 |
| HCP (B3) | no | 0.949 | 0.956 | 0.933 | 0.949 | 1.441 | 0.559 | 0.127 | 0.000 |
| HCP (B3) | yes | 0.953 | 0.957 | 0.947 | 0.948 | 1.447 | 0.553 | 0.130 | 0.000 |
| Subsample once (B3) | no | 0.947 | 0.955 | 0.931 | 0.948 | 1.437 | 0.563 | 0.131 | 0.000 |
| Subsample once (B3) | yes | 0.952 | 0.955 | 0.946 | 0.947 | 1.441 | 0.559 | 0.133 | 0.000 |
| Repeated subs. (B3) | no | 0.949 | 0.956 | 0.934 | 0.950 | 1.441 | 0.559 | 0.127 | 0.000 |
| Repeated subs. (B3) | yes | 0.953 | 0.957 | 0.947 | 0.948 | 1.447 | 0.553 | 0.130 | 0.000 |
| GWCP (B4) | no | 0.949 | 0.956 | 0.934 | 0.949 | 1.442 | 0.558 | 0.128 | 0.000 |

*All methods at $\alpha = 0.05$, continued*

| Method | cc | Cov | var.-w. | $\mathrm{Cov}_1$ | $\mathrm{Cov}_0$ | size | single. | under. | abst. |
|---|---|---|---|---|---|---|---|---|---|
| GWCP (B4) | yes | 0.954 | 0.957 | 0.948 | 0.949 | 1.451 | 0.549 | 0.128 | 0.000 |
| ProtCOVER | no | 0.954 | 0.960 | 0.942 | 0.953 | 1.469 | 0.531 | 0.111 | 0.000 |
| ProtCOVER | yes | 0.959 | 0.961 | 0.956 | 0.953 | 1.478 | 0.522 | 0.109 | 0.000 |
| ProtCOVER-D | no | 0.955 | 0.952 | 0.929 | 0.960 | 1.425 | 0.575 | 0.117 | 0.000 |
| ProtCOVER-D | yes | 0.958 | 0.960 | 0.955 | 0.955 | 1.484 | 0.516 | 0.117 | 0.000 |
| ProtCOVER-W | no | 0.953 | 0.960 | 0.939 | 0.952 | 1.462 | 0.538 | 0.113 | 0.000 |
| ProtCOVER-W | yes | 0.957 | 0.960 | 0.954 | 0.951 | 1.474 | 0.526 | 0.112 | 0.000 |
| (i)+(iii) gate | no | 0.949 | 0.956 | 0.933 | 0.949 | 1.441 | 0.559 | 0.127 | 0.000 |
| (i)+(iii) gate | yes | 0.953 | 0.957 | 0.947 | 0.948 | 1.447 | 0.553 | 0.130 | 0.000 |
| ProtCOVER-W + gate | no | 0.953 | 0.960 | 0.939 | 0.952 | 1.462 | 0.538 | 0.113 | 0.000 |
| ProtCOVER-W + gate | yes | 0.957 | 0.960 | 0.954 | 0.951 | 1.474 | 0.526 | 0.112 | 0.000 |
| Support-only (B5) | no | 0.953 | 0.960 | 0.939 | 0.952 | 1.462 | 0.538 | 0.113 | 0.000 |
| Support-only (B5) | yes | 0.957 | 0.960 | 0.954 | 0.951 | 1.474 | 0.526 | 0.112 | 0.000 |
| Random abst. (B5) | no | 0.953 | 0.960 | 0.939 | 0.952 | 1.462 | 0.538 | 0.113 | 0.000 |
| Random abst. (B5) | yes | 0.957 | 0.960 | 0.954 | 0.951 | 1.474 | 0.526 | 0.112 | 0.000 |
| *S2*, $\beta = 1$ | | | | | | | | | |
| Split (B1/B2) | no | 0.937 | 0.945 | 0.913 | 0.937 | 1.364 | 0.636 | 0.167 | 0.000 |
| Split (B1/B2) | yes | 0.939 | 0.944 | 0.902 | 0.945 | 1.361 | 0.639 | 0.165 | 0.000 |
| HCP (B3) | no | 0.947 | 0.955 | 0.929 | 0.949 | 1.439 | 0.561 | 0.133 | 0.000 |
| HCP (B3) | yes | 0.952 | 0.955 | 0.941 | 0.948 | 1.442 | 0.558 | 0.134 | 0.000 |
| Subsample once (B3) | no | 0.948 | 0.955 | 0.931 | 0.948 | 1.445 | 0.555 | 0.134 | 0.000 |
| Subsample once (B3) | yes | 0.950 | 0.953 | 0.939 | 0.946 | 1.434 | 0.566 | 0.142 | 0.000 |
| Repeated subs. (B3) | no | 0.947 | 0.955 | 0.930 | 0.949 | 1.440 | 0.560 | 0.134 | 0.000 |
| Repeated subs. (B3) | yes | 0.952 | 0.955 | 0.941 | 0.948 | 1.441 | 0.559 | 0.135 | 0.000 |
| GWCP (B4) | no | 0.948 | 0.956 | 0.932 | 0.949 | 1.448 | 0.552 | 0.129 | 0.000 |
| GWCP (B4) | yes | 0.954 | 0.957 | 0.946 | 0.950 | 1.455 | 0.545 | 0.129 | 0.000 |
| ProtCOVER | no | 0.958 | 0.963 | 0.947 | 0.958 | 1.502 | 0.498 | 0.102 | 0.000 |
| ProtCOVER | yes | 0.961 | 0.962 | 0.956 | 0.957 | 1.499 | 0.501 | 0.104 | 0.000 |
| ProtCOVER-D | no | 0.955 | 0.952 | 0.925 | 0.962 | 1.429 | 0.571 | 0.118 | 0.000 |
| ProtCOVER-D | yes | 0.959 | 0.961 | 0.955 | 0.958 | 1.496 | 0.504 | 0.113 | 0.000 |
| ProtCOVER-W | no | 0.955 | 0.961 | 0.940 | 0.955 | 1.486 | 0.514 | 0.107 | 0.000 |
| ProtCOVER-W | yes | 0.959 | 0.961 | 0.953 | 0.955 | 1.485 | 0.515 | 0.106 | 0.000 |
| (i)+(iii) gate | no | 0.947 | 0.955 | 0.929 | 0.949 | 1.439 | 0.561 | 0.133 | 0.000 |
| (i)+(iii) gate | yes | 0.952 | 0.955 | 0.941 | 0.948 | 1.442 | 0.558 | 0.134 | 0.000 |
| ProtCOVER-W + gate | no | 0.955 | 0.961 | 0.940 | 0.955 | 1.486 | 0.514 | 0.107 | 0.000 |
| ProtCOVER-W + gate | yes | 0.959 | 0.961 | 0.953 | 0.955 | 1.485 | 0.515 | 0.106 | 0.000 |
| Support-only (B5) | no | 0.955 | 0.961 | 0.940 | 0.955 | 1.486 | 0.514 | 0.107 | 0.000 |
| Support-only (B5) | yes | 0.959 | 0.961 | 0.953 | 0.955 | 1.485 | 0.515 | 0.106 | 0.000 |
| Random abst. (B5) | no | 0.955 | 0.961 | 0.940 | 0.955 | 1.486 | 0.514 | 0.107 | 0.000 |
| Random abst. (B5) | yes | 0.959 | 0.961 | 0.953 | 0.955 | 1.485 | 0.515 | 0.106 | 0.000 |
| *S2*, $\beta = 2$ | | | | | | | | | |
| Split (B1/B2) | no | 0.935 | 0.943 | 0.908 | 0.936 | 1.361 | 0.639 | 0.175 | 0.000 |
| Split (B1/B2) | yes | 0.937 | 0.942 | 0.895 | 0.944 | 1.357 | 0.643 | 0.173 | 0.000 |
| HCP (B3) | no | 0.946 | 0.953 | 0.926 | 0.947 | 1.435 | 0.565 | 0.140 | 0.000 |
| HCP (B3) | yes | 0.948 | 0.952 | 0.934 | 0.945 | 1.431 | 0.569 | 0.147 | 0.000 |
| Subsample once (B3) | no | 0.944 | 0.950 | 0.920 | 0.947 | 1.418 | 0.582 | 0.148 | 0.000 |
| Subsample once (B3) | yes | 0.947 | 0.951 | 0.934 | 0.944 | 1.428 | 0.572 | 0.151 | 0.000 |
| Repeated subs. (B3) | no | 0.946 | 0.953 | 0.926 | 0.947 | 1.435 | 0.565 | 0.140 | 0.000 |
| Repeated subs. (B3) | yes | 0.948 | 0.952 | 0.934 | 0.945 | 1.431 | 0.569 | 0.147 | 0.000 |

*All methods at $\alpha = 0.05$, continued*

| Method | cc | Cov | var.-w. | $Cov_1$ | $Cov_0$ | size | single. | under. | abst. |
|---|---|---|---|---|---|---|---|---|---|
| GWCP (B4) | no | 0.947 | 0.954 | 0.929 | 0.948 | 1.452 | 0.548 | 0.139 | 0.000 |
| GWCP (B4) | yes | 0.948 | 0.951 | 0.943 | 0.942 | 1.448 | 0.552 | 0.157 | 0.000 |
| ProtCOVER | no | 0.969 | 0.972 | 0.962 | 0.968 | 1.625 | 0.375 | 0.074 | 0.015 |
| ProtCOVER | yes | 0.971 | 0.972 | 0.969 | 0.967 | 1.621 | 0.379 | 0.079 | 0.015 |
| ProtCOVER-D | no | 0.955 | 0.953 | 0.924 | 0.961 | 1.437 | 0.563 | 0.121 | 0.000 |
| ProtCOVER-D | yes | 0.959 | 0.960 | 0.955 | 0.956 | 1.513 | 0.487 | 0.117 | 0.000 |
| ProtCOVER-W | no | 0.962 | 0.967 | 0.952 | 0.961 | 1.549 | 0.451 | 0.090 | 0.000 |
| ProtCOVER-W | yes | 0.964 | 0.966 | 0.962 | 0.959 | 1.541 | 0.459 | 0.094 | 0.000 |
| (i)+(iii) gate | no | 0.946 | 0.953 | 0.926 | 0.947 | 1.435 | 0.565 | 0.140 | 0.000 |
| (i)+(iii) gate | yes | 0.948 | 0.952 | 0.934 | 0.945 | 1.431 | 0.569 | 0.147 | 0.000 |
| ProtCOVER-W + gate | no | 0.962 | 0.967 | 0.952 | 0.961 | 1.549 | 0.451 | 0.090 | 0.000 |
| ProtCOVER-W + gate | yes | 0.964 | 0.966 | 0.962 | 0.959 | 1.543 | 0.457 | 0.094 | 0.007 |
| Support-only (B5) | no | 0.962 | 0.967 | 0.952 | 0.961 | 1.549 | 0.451 | 0.090 | 0.000 |
| Support-only (B5) | yes | 0.964 | 0.966 | 0.962 | 0.959 | 1.543 | 0.457 | 0.094 | 0.007 |
| Random abst. (B5) | no | 0.962 | 0.967 | 0.952 | 0.961 | 1.549 | 0.451 | 0.090 | 0.000 |
| Random abst. (B5) | yes | 0.964 | 0.966 | 0.962 | 0.959 | 1.544 | 0.456 | 0.094 | 0.007 |
| *S3 (orphan families)* | | | | | | | | | |
| Split (B1/B2) | no | 0.938 | 0.946 | 0.924 | 0.935 | 1.371 | 0.629 | 0.154 | 0.000 |
| Split (B1/B2) | yes | 0.941 | 0.946 | 0.918 | 0.942 | 1.371 | 0.629 | 0.149 | 0.000 |
| HCP (B3) | no | 0.950 | 0.956 | 0.938 | 0.950 | 1.451 | 0.549 | 0.116 | 0.000 |
| HCP (B3) | yes | 0.956 | 0.959 | 0.954 | 0.949 | 1.459 | 0.541 | 0.117 | 0.000 |
| Subsample once (B3) | no | 0.948 | 0.954 | 0.936 | 0.947 | 1.442 | 0.558 | 0.128 | 0.000 |
| Subsample once (B3) | yes | 0.955 | 0.957 | 0.953 | 0.948 | 1.454 | 0.546 | 0.118 | 0.000 |
| Repeated subs. (B3) | no | 0.950 | 0.955 | 0.937 | 0.950 | 1.449 | 0.551 | 0.116 | 0.000 |
| Repeated subs. (B3) | yes | 0.956 | 0.959 | 0.955 | 0.950 | 1.459 | 0.541 | 0.116 | 0.000 |
| GWCP (B4) | no | 0.951 | 0.956 | 0.938 | 0.953 | 1.461 | 0.539 | 0.117 | 0.000 |
| GWCP (B4) | yes | 0.956 | 0.957 | 0.949 | 0.952 | 1.448 | 0.552 | 0.118 | 0.000 |
| ProtCOVER | no | 0.956 | 0.960 | 0.943 | 0.958 | 1.486 | 0.514 | 0.101 | 0.000 |
| ProtCOVER | yes | 0.960 | 0.961 | 0.953 | 0.958 | 1.477 | 0.523 | 0.099 | 0.000 |
| ProtCOVER-D | no | 0.958 | 0.956 | 0.935 | 0.963 | 1.438 | 0.562 | 0.109 | 0.000 |
| ProtCOVER-D | yes | 0.959 | 0.959 | 0.955 | 0.958 | 1.461 | 0.539 | 0.111 | 0.000 |
| ProtCOVER-W | no | 0.956 | 0.959 | 0.942 | 0.958 | 1.485 | 0.515 | 0.104 | 0.000 |
| ProtCOVER-W | yes | 0.960 | 0.961 | 0.951 | 0.959 | 1.473 | 0.527 | 0.100 | 0.000 |
| (i)+(iii) gate | no | 0.950 | 0.956 | 0.938 | 0.950 | 1.451 | 0.549 | 0.116 | 0.000 |
| (i)+(iii) gate | yes | 0.956 | 0.959 | 0.955 | 0.950 | 1.460 | 0.540 | 0.116 | 0.003 |
| ProtCOVER-W + gate | no | 0.956 | 0.959 | 0.942 | 0.958 | 1.485 | 0.515 | 0.104 | 0.000 |
| ProtCOVER-W + gate | yes | 0.960 | 0.961 | 0.951 | 0.959 | 1.473 | 0.527 | 0.100 | 0.001 |
| Support-only (B5) | no | 0.956 | 0.959 | 0.942 | 0.958 | 1.485 | 0.515 | 0.104 | 0.000 |
| Support-only (B5) | yes | 0.960 | 0.961 | 0.951 | 0.959 | 1.473 | 0.527 | 0.100 | 0.001 |
| Random abst. (B5) | no | 0.956 | 0.959 | 0.942 | 0.958 | 1.485 | 0.515 | 0.104 | 0.000 |
| Random abst. (B5) | yes | 0.960 | 0.961 | 0.951 | 0.959 | 1.473 | 0.527 | 0.100 | 0.001 |
| *S4 (held-out regions)* | | | | | | | | | |
| Split (B1/B2) | no | 0.939 | 0.947 | 0.925 | 0.938 | 1.377 | 0.623 | 0.152 | 0.000 |
| Split (B1/B2) | yes | 0.941 | 0.947 | 0.922 | 0.942 | 1.375 | 0.625 | 0.145 | 0.000 |
| HCP (B3) | no | 0.949 | 0.956 | 0.936 | 0.951 | 1.444 | 0.556 | 0.122 | 0.000 |
| HCP (B3) | yes | 0.954 | 0.958 | 0.951 | 0.948 | 1.446 | 0.554 | 0.118 | 0.000 |
| Subsample once (B3) | no | 0.948 | 0.955 | 0.936 | 0.949 | 1.443 | 0.557 | 0.125 | 0.000 |
| Subsample once (B3) | yes | 0.953 | 0.958 | 0.951 | 0.947 | 1.442 | 0.558 | 0.122 | 0.000 |
| Repeated subs. (B3) | no | 0.949 | 0.956 | 0.937 | 0.950 | 1.445 | 0.555 | 0.122 | 0.000 |

*All methods at $\alpha = 0.05$, continued*

| Method | cc | Cov | var.-w. | $\text{Cov}_1$ | $\text{Cov}_0$ | size | single. | under. | abst. |
|---|---|---|---|---|---|---|---|---|---|
| Repeated subs. (B3) | yes | 0.954 | 0.958 | 0.951 | 0.948 | 1.445 | 0.555 | 0.119 | 0.000 |
| GWCP (B4) | no | 0.949 | 0.955 | 0.936 | 0.950 | 1.440 | 0.560 | 0.127 | 0.000 |
| GWCP (B4) | yes | 0.953 | 0.956 | 0.948 | 0.948 | 1.436 | 0.564 | 0.125 | 0.000 |
| ProtCOVER | no | 0.956 | 0.961 | 0.944 | 0.958 | 1.495 | 0.505 | 0.107 | 0.028 |
| ProtCOVER | yes | 0.960 | 0.962 | 0.958 | 0.955 | 1.495 | 0.505 | 0.108 | 0.028 |
| ProtCOVER-D | no | 0.958 | 0.956 | 0.936 | 0.964 | 1.458 | 0.542 | 0.106 | 0.000 |
| ProtCOVER-D | yes | 0.958 | 0.960 | 0.954 | 0.957 | 1.481 | 0.519 | 0.117 | 0.000 |
| ProtCOVER-W | no | 0.954 | 0.959 | 0.941 | 0.957 | 1.485 | 0.515 | 0.111 | 0.000 |
| ProtCOVER-W | yes | 0.956 | 0.957 | 0.949 | 0.953 | 1.465 | 0.535 | 0.122 | 0.000 |
| (i)+(iii) gate | no | 0.949 | 0.956 | 0.936 | 0.951 | 1.444 | 0.556 | 0.122 | 0.000 |
| (i)+(iii) gate | yes | 0.954 | 0.958 | 0.952 | 0.948 | 1.448 | 0.552 | 0.118 | 0.002 |
| ProtCOVER-W + gate | no | 0.954 | 0.959 | 0.941 | 0.957 | 1.485 | 0.515 | 0.111 | 0.000 |
| ProtCOVER-W + gate | yes | 0.956 | 0.957 | 0.949 | 0.953 | 1.465 | 0.535 | 0.122 | 0.002 |
| Support-only (B5) | no | 0.954 | 0.959 | 0.941 | 0.957 | 1.485 | 0.515 | 0.111 | 0.000 |
| Support-only (B5) | yes | 0.956 | 0.957 | 0.949 | 0.953 | 1.465 | 0.535 | 0.122 | 0.002 |
| Random abst. (B5) | no | 0.954 | 0.959 | 0.941 | 0.957 | 1.485 | 0.515 | 0.111 | 0.000 |
| Random abst. (B5) | yes | 0.956 | 0.957 | 0.949 | 0.953 | 1.465 | 0.535 | 0.122 | 0.002 |

Table 13: All methods at $\alpha = 0.2$, mean over predictors and seeds.

| Method | cc | Cov | var.-w. | $\text{Cov}_1$ | $\text{Cov}_0$ | size | single. | under. | abst. |
|---|---|---|---|---|---|---|---|---|---|
| *S1 (exchangeable)* | | | | | | | | | |
| Split (B1/B2) | no | 0.797 | 0.794 | 0.753 | 0.804 | 0.954 | 0.934 | 0.280 | 0.000 |
| Split (B1/B2) | yes | 0.791 | 0.793 | 0.758 | 0.792 | 0.947 | 0.938 | 0.290 | 0.000 |
| HCP (B3) | no | 0.799 | 0.797 | 0.755 | 0.807 | 0.958 | 0.943 | 0.272 | 0.000 |
| HCP (B3) | yes | 0.812 | 0.813 | 0.800 | 0.800 | 0.994 | 0.939 | 0.246 | 0.000 |
| Subsample once (B3) | no | 0.799 | 0.797 | 0.753 | 0.808 | 0.959 | 0.938 | 0.273 | 0.000 |
| Subsample once (B3) | yes | 0.812 | 0.812 | 0.799 | 0.801 | 0.994 | 0.937 | 0.251 | 0.000 |
| Repeated subs. (B3) | no | 0.799 | 0.797 | 0.755 | 0.807 | 0.958 | 0.943 | 0.271 | 0.000 |
| Repeated subs. (B3) | yes | 0.812 | 0.813 | 0.800 | 0.800 | 0.994 | 0.939 | 0.245 | 0.000 |
| GWCP (B4) | no | 0.798 | 0.795 | 0.754 | 0.805 | 0.955 | 0.942 | 0.276 | 0.000 |
| GWCP (B4) | yes | 0.811 | 0.812 | 0.800 | 0.799 | 0.992 | 0.938 | 0.249 | 0.000 |
| ProtCOVER | no | 0.802 | 0.796 | 0.757 | 0.810 | 0.956 | 0.941 | 0.269 | 0.000 |
| ProtCOVER | yes | 0.812 | 0.813 | 0.802 | 0.803 | 0.993 | 0.941 | 0.251 | 0.000 |
| ProtCOVER-D | no | 0.803 | 0.793 | 0.755 | 0.815 | 0.946 | 0.929 | 0.273 | 0.000 |
| ProtCOVER-D | yes | 0.808 | 0.810 | 0.806 | 0.803 | 0.981 | 0.947 | 0.264 | 0.000 |
| ProtCOVER-W | no | 0.800 | 0.798 | 0.756 | 0.807 | 0.959 | 0.945 | 0.270 | 0.000 |
| ProtCOVER-W | yes | 0.812 | 0.813 | 0.799 | 0.801 | 0.994 | 0.937 | 0.247 | 0.000 |
| (i)+(iii) gate | no | 0.799 | 0.797 | 0.755 | 0.807 | 0.958 | 0.943 | 0.272 | 0.000 |
| (i)+(iii) gate | yes | 0.812 | 0.813 | 0.800 | 0.800 | 0.994 | 0.939 | 0.246 | 0.000 |
| ProtCOVER-W + gate | no | 0.800 | 0.798 | 0.756 | 0.808 | 0.959 | 0.944 | 0.269 | 0.000 |
| ProtCOVER-W + gate | yes | 0.812 | 0.813 | 0.799 | 0.802 | 0.995 | 0.936 | 0.247 | 0.002 |
| Support-only (B5) | no | 0.800 | 0.798 | 0.756 | 0.807 | 0.959 | 0.944 | 0.270 | 0.000 |
| Support-only (B5) | yes | 0.812 | 0.813 | 0.799 | 0.802 | 0.995 | 0.936 | 0.247 | 0.002 |
| Random abst. (B5) | no | 0.800 | 0.798 | 0.756 | 0.807 | 0.959 | 0.944 | 0.270 | 0.000 |
| Random abst. (B5) | yes | 0.812 | 0.813 | 0.799 | 0.802 | 0.996 | 0.936 | 0.247 | 0.002 |
| *S2, $\beta = 0.5$* | | | | | | | | | |
| Split (B1/B2) | no | 0.792 | 0.791 | 0.741 | 0.802 | 0.950 | 0.928 | 0.294 | 0.000 |

*All methods at $\alpha = 0.2$, continued*

| Method | cc | Cov | var.-w. | $\text{Cov}_1$ | $\text{Cov}_0$ | size | single. | under. | abst. |
|---|---|---|---|---|---|---|---|---|---|
| Split (B1/B2) | yes | 0.787 | 0.789 | 0.741 | 0.794 | 0.944 | 0.933 | 0.304 | 0.000 |
| HCP (B3) | no | 0.794 | 0.792 | 0.742 | 0.804 | 0.952 | 0.941 | 0.289 | 0.000 |
| HCP (B3) | yes | 0.805 | 0.804 | 0.783 | 0.798 | 0.982 | 0.937 | 0.267 | 0.000 |
| Subsample once (B3) | no | 0.791 | 0.789 | 0.737 | 0.803 | 0.946 | 0.937 | 0.298 | 0.000 |
| Subsample once (B3) | yes | 0.804 | 0.803 | 0.783 | 0.797 | 0.982 | 0.937 | 0.271 | 0.000 |
| Repeated subs. (B3) | no | 0.794 | 0.792 | 0.743 | 0.804 | 0.952 | 0.942 | 0.289 | 0.000 |
| Repeated subs. (B3) | yes | 0.805 | 0.804 | 0.782 | 0.798 | 0.981 | 0.937 | 0.268 | 0.000 |
| GWCP (B4) | no | 0.795 | 0.793 | 0.743 | 0.805 | 0.954 | 0.943 | 0.285 | 0.000 |
| GWCP (B4) | yes | 0.809 | 0.811 | 0.794 | 0.798 | 0.995 | 0.937 | 0.258 | 0.000 |
| ProtCOVER | no | 0.798 | 0.796 | 0.749 | 0.808 | 0.960 | 0.944 | 0.278 | 0.000 |
| ProtCOVER | yes | 0.810 | 0.807 | 0.799 | 0.801 | 0.997 | 0.931 | 0.259 | 0.000 |
| ProtCOVER-D | no | 0.801 | 0.792 | 0.745 | 0.814 | 0.949 | 0.932 | 0.280 | 0.000 |
| ProtCOVER-D | yes | 0.807 | 0.810 | 0.801 | 0.804 | 0.988 | 0.949 | 0.266 | 0.000 |
| ProtCOVER-W | no | 0.797 | 0.796 | 0.746 | 0.807 | 0.959 | 0.946 | 0.280 | 0.000 |
| ProtCOVER-W | yes | 0.813 | 0.815 | 0.801 | 0.801 | 1.004 | 0.939 | 0.251 | 0.000 |
| (i)+(iii) gate | no | 0.794 | 0.792 | 0.742 | 0.804 | 0.952 | 0.941 | 0.289 | 0.000 |
| (i)+(iii) gate | yes | 0.805 | 0.804 | 0.783 | 0.799 | 0.982 | 0.937 | 0.267 | 0.000 |
| ProtCOVER-W + gate | no | 0.797 | 0.796 | 0.746 | 0.807 | 0.959 | 0.945 | 0.279 | 0.000 |
| ProtCOVER-W + gate | yes | 0.813 | 0.815 | 0.801 | 0.801 | 1.005 | 0.938 | 0.250 | 0.001 |
| Support-only (B5) | no | 0.797 | 0.796 | 0.746 | 0.807 | 0.959 | 0.945 | 0.279 | 0.000 |
| Support-only (B5) | yes | 0.813 | 0.815 | 0.801 | 0.801 | 1.005 | 0.937 | 0.250 | 0.001 |
| Random abst. (B5) | no | 0.797 | 0.796 | 0.746 | 0.807 | 0.959 | 0.945 | 0.280 | 0.000 |
| Random abst. (B5) | yes | 0.813 | 0.815 | 0.801 | 0.801 | 1.005 | 0.937 | 0.250 | 0.001 |
| *S2*, $\beta = 1$ | | | | | | | | | |
| Split (B1/B2) | no | 0.790 | 0.788 | 0.732 | 0.801 | 0.947 | 0.924 | 0.307 | 0.000 |
| Split (B1/B2) | yes | 0.784 | 0.785 | 0.728 | 0.795 | 0.942 | 0.929 | 0.313 | 0.000 |
| HCP (B3) | no | 0.791 | 0.789 | 0.733 | 0.803 | 0.949 | 0.938 | 0.302 | 0.000 |
| HCP (B3) | yes | 0.800 | 0.798 | 0.766 | 0.799 | 0.973 | 0.933 | 0.285 | 0.000 |
| Subsample once (B3) | no | 0.786 | 0.784 | 0.726 | 0.798 | 0.940 | 0.930 | 0.317 | 0.000 |
| Subsample once (B3) | yes | 0.799 | 0.797 | 0.764 | 0.798 | 0.971 | 0.933 | 0.287 | 0.000 |
| Repeated subs. (B3) | no | 0.791 | 0.789 | 0.733 | 0.803 | 0.949 | 0.938 | 0.301 | 0.000 |
| Repeated subs. (B3) | yes | 0.800 | 0.797 | 0.766 | 0.798 | 0.973 | 0.933 | 0.286 | 0.000 |
| GWCP (B4) | no | 0.793 | 0.792 | 0.735 | 0.805 | 0.953 | 0.941 | 0.294 | 0.000 |
| GWCP (B4) | yes | 0.807 | 0.808 | 0.787 | 0.798 | 0.995 | 0.935 | 0.268 | 0.000 |
| ProtCOVER | no | 0.804 | 0.801 | 0.751 | 0.815 | 0.976 | 0.935 | 0.270 | 0.000 |
| ProtCOVER | yes | 0.818 | 0.813 | 0.802 | 0.810 | 1.020 | 0.912 | 0.248 | 0.000 |
| ProtCOVER-D | no | 0.799 | 0.792 | 0.738 | 0.814 | 0.952 | 0.933 | 0.286 | 0.000 |
| ProtCOVER-D | yes | 0.808 | 0.810 | 0.794 | 0.807 | 0.992 | 0.951 | 0.268 | 0.000 |
| ProtCOVER-W | no | 0.800 | 0.798 | 0.745 | 0.811 | 0.966 | 0.949 | 0.275 | 0.000 |
| ProtCOVER-W | yes | 0.816 | 0.817 | 0.800 | 0.805 | 1.015 | 0.938 | 0.245 | 0.000 |
| (i)+(iii) gate | no | 0.791 | 0.789 | 0.733 | 0.803 | 0.949 | 0.938 | 0.302 | 0.000 |
| (i)+(iii) gate | yes | 0.800 | 0.798 | 0.766 | 0.799 | 0.974 | 0.933 | 0.285 | 0.000 |
| ProtCOVER-W + gate | no | 0.800 | 0.798 | 0.745 | 0.811 | 0.966 | 0.949 | 0.274 | 0.000 |
| ProtCOVER-W + gate | yes | 0.816 | 0.818 | 0.801 | 0.805 | 1.016 | 0.937 | 0.244 | 0.001 |
| Support-only (B5) | no | 0.800 | 0.798 | 0.745 | 0.811 | 0.966 | 0.949 | 0.274 | 0.000 |
| Support-only (B5) | yes | 0.816 | 0.818 | 0.801 | 0.805 | 1.016 | 0.937 | 0.244 | 0.001 |
| Random abst. (B5) | no | 0.800 | 0.798 | 0.745 | 0.811 | 0.966 | 0.949 | 0.274 | 0.000 |
| Random abst. (B5) | yes | 0.816 | 0.818 | 0.801 | 0.805 | 1.016 | 0.937 | 0.244 | 0.001 |

*All methods at $\alpha = 0.2$, continued*

| Method | cc | Cov | var.-w. | $Cov_1$ | $Cov_0$ | size | single. | under. | abst. |
|---|---|---|---|---|---|---|---|---|---|
| *S2, $\beta = 2$* | | | | | | | | | |
| Split (B1/B2) | no | 0.789 | 0.787 | 0.726 | 0.801 | 0.947 | 0.922 | 0.309 | 0.000 |
| Split (B1/B2) | yes | 0.782 | 0.784 | 0.722 | 0.793 | 0.940 | 0.926 | 0.318 | 0.000 |
| HCP (B3) | no | 0.790 | 0.788 | 0.727 | 0.802 | 0.947 | 0.936 | 0.305 | 0.000 |
| HCP (B3) | yes | 0.795 | 0.793 | 0.757 | 0.795 | 0.966 | 0.931 | 0.300 | 0.000 |
| Subsample once (B3) | no | 0.789 | 0.786 | 0.723 | 0.802 | 0.944 | 0.935 | 0.308 | 0.000 |
| Subsample once (B3) | yes | 0.795 | 0.792 | 0.755 | 0.795 | 0.966 | 0.930 | 0.301 | 0.000 |
| Repeated subs. (B3) | no | 0.790 | 0.788 | 0.727 | 0.802 | 0.947 | 0.936 | 0.305 | 0.000 |
| Repeated subs. (B3) | yes | 0.795 | 0.793 | 0.757 | 0.795 | 0.966 | 0.931 | 0.300 | 0.000 |
| GWCP (B4) | no | 0.796 | 0.794 | 0.733 | 0.808 | 0.959 | 0.935 | 0.291 | 0.000 |
| GWCP (B4) | yes | 0.803 | 0.805 | 0.779 | 0.794 | 0.992 | 0.925 | 0.283 | 0.000 |
| ProtCOVER | no | 0.829 | 0.826 | 0.785 | 0.836 | 1.064 | 0.862 | 0.224 | 0.000 |
| ProtCOVER | yes | 0.835 | 0.830 | 0.832 | 0.821 | 1.114 | 0.826 | 0.221 | 0.000 |
| ProtCOVER-D | no | 0.799 | 0.795 | 0.733 | 0.814 | 0.959 | 0.936 | 0.285 | 0.000 |
| ProtCOVER-D | yes | 0.805 | 0.808 | 0.783 | 0.807 | 0.998 | 0.947 | 0.274 | 0.000 |
| ProtCOVER-W | no | 0.806 | 0.804 | 0.748 | 0.817 | 0.978 | 0.942 | 0.263 | 0.000 |
| ProtCOVER-W | yes | 0.815 | 0.815 | 0.796 | 0.805 | 1.015 | 0.928 | 0.253 | 0.000 |
| (i)+(iii) gate | no | 0.790 | 0.788 | 0.727 | 0.802 | 0.948 | 0.936 | 0.305 | 0.000 |
| (i)+(iii) gate | yes | 0.796 | 0.793 | 0.757 | 0.795 | 0.966 | 0.931 | 0.300 | 0.000 |
| ProtCOVER-W + gate | no | 0.812 | 0.810 | 0.755 | 0.822 | 1.010 | 0.912 | 0.254 | 0.030 |
| ProtCOVER-W + gate | yes | 0.820 | 0.820 | 0.801 | 0.810 | 1.040 | 0.903 | 0.247 | 0.030 |
| Support-only (B5) | no | 0.812 | 0.809 | 0.754 | 0.822 | 1.005 | 0.918 | 0.256 | 0.030 |
| Support-only (B5) | yes | 0.820 | 0.819 | 0.800 | 0.810 | 1.040 | 0.903 | 0.248 | 0.030 |
| Random abst. (B5) | no | 0.812 | 0.810 | 0.755 | 0.822 | 1.010 | 0.914 | 0.254 | 0.030 |
| Random abst. (B5) | yes | 0.820 | 0.820 | 0.800 | 0.810 | 1.041 | 0.901 | 0.247 | 0.030 |
| *S3 (orphan families)* | | | | | | | | | |
| Split (B1/B2) | no | 0.794 | 0.793 | 0.750 | 0.801 | 0.952 | 0.927 | 0.295 | 0.000 |
| Split (B1/B2) | yes | 0.787 | 0.789 | 0.754 | 0.787 | 0.943 | 0.934 | 0.308 | 0.000 |
| HCP (B3) | no | 0.797 | 0.796 | 0.753 | 0.804 | 0.956 | 0.938 | 0.284 | 0.000 |
| HCP (B3) | yes | 0.811 | 0.813 | 0.803 | 0.795 | 0.989 | 0.940 | 0.259 | 0.000 |
| Subsample once (B3) | no | 0.798 | 0.796 | 0.750 | 0.809 | 0.959 | 0.938 | 0.282 | 0.000 |
| Subsample once (B3) | yes | 0.811 | 0.813 | 0.804 | 0.795 | 0.991 | 0.940 | 0.259 | 0.000 |
| Repeated subs. (B3) | no | 0.797 | 0.796 | 0.753 | 0.804 | 0.956 | 0.938 | 0.285 | 0.000 |
| Repeated subs. (B3) | yes | 0.811 | 0.813 | 0.803 | 0.795 | 0.989 | 0.940 | 0.258 | 0.000 |
| GWCP (B4) | no | 0.803 | 0.802 | 0.760 | 0.810 | 0.967 | 0.945 | 0.266 | 0.000 |
| GWCP (B4) | yes | 0.814 | 0.814 | 0.800 | 0.804 | 0.993 | 0.941 | 0.250 | 0.000 |
| ProtCOVER | no | 0.806 | 0.805 | 0.764 | 0.813 | 0.974 | 0.942 | 0.260 | 0.000 |
| ProtCOVER | yes | 0.818 | 0.817 | 0.798 | 0.812 | 1.001 | 0.941 | 0.244 | 0.000 |
| ProtCOVER-D | no | 0.810 | 0.803 | 0.762 | 0.820 | 0.967 | 0.940 | 0.264 | 0.000 |
| ProtCOVER-D | yes | 0.813 | 0.811 | 0.806 | 0.811 | 0.993 | 0.951 | 0.254 | 0.000 |
| ProtCOVER-W | no | 0.807 | 0.806 | 0.764 | 0.814 | 0.975 | 0.943 | 0.258 | 0.000 |
| ProtCOVER-W | yes | 0.819 | 0.817 | 0.796 | 0.813 | 1.000 | 0.942 | 0.242 | 0.000 |
| (i)+(iii) gate | no | 0.797 | 0.796 | 0.753 | 0.804 | 0.956 | 0.938 | 0.284 | 0.000 |
| (i)+(iii) gate | yes | 0.811 | 0.813 | 0.804 | 0.795 | 0.990 | 0.939 | 0.258 | 0.002 |
| ProtCOVER-W + gate | no | 0.807 | 0.806 | 0.765 | 0.814 | 0.977 | 0.942 | 0.257 | 0.002 |
| ProtCOVER-W + gate | yes | 0.819 | 0.817 | 0.797 | 0.814 | 1.003 | 0.939 | 0.241 | 0.004 |
| Support-only (B5) | no | 0.807 | 0.806 | 0.765 | 0.814 | 0.977 | 0.942 | 0.257 | 0.002 |
| Support-only (B5) | yes | 0.819 | 0.817 | 0.797 | 0.814 | 1.003 | 0.939 | 0.240 | 0.004 |
| Random abst. (B5) | no | 0.807 | 0.806 | 0.765 | 0.814 | 0.977 | 0.942 | 0.257 | 0.002 |

*All methods at $\alpha = 0.2$, continued*

| Method | cc | Cov | var.-w. | $\text{Cov}_1$ | $\text{Cov}_0$ | size | single. | under. | abst. |
|---|---|---|---|---|---|---|---|---|---|
| Random abst. (B5) | yes | 0.819 | 0.817 | 0.797 | 0.814 | 1.004 | 0.938 | 0.240 | 0.004 |
| *S4 (held-out regions)* | | | | | | | | | |
| Split (B1/B2) | no | 0.797 | 0.792 | 0.754 | 0.811 | 0.959 | 0.930 | 0.283 | 0.000 |
| Split (B1/B2) | yes | 0.791 | 0.793 | 0.763 | 0.792 | 0.951 | 0.937 | 0.294 | 0.000 |
| HCP (B3) | no | 0.796 | 0.790 | 0.753 | 0.810 | 0.955 | 0.941 | 0.280 | 0.000 |
| HCP (B3) | yes | 0.811 | 0.812 | 0.803 | 0.799 | 0.993 | 0.937 | 0.250 | 0.000 |
| Subsample once (B3) | no | 0.796 | 0.790 | 0.752 | 0.810 | 0.956 | 0.941 | 0.280 | 0.000 |
| Subsample once (B3) | yes | 0.810 | 0.812 | 0.804 | 0.797 | 0.992 | 0.936 | 0.252 | 0.000 |
| Repeated subs. (B3) | no | 0.796 | 0.790 | 0.752 | 0.810 | 0.955 | 0.941 | 0.280 | 0.000 |
| Repeated subs. (B3) | yes | 0.811 | 0.812 | 0.803 | 0.799 | 0.993 | 0.937 | 0.251 | 0.000 |
| GWCP (B4) | no | 0.798 | 0.792 | 0.754 | 0.812 | 0.959 | 0.944 | 0.275 | 0.000 |
| GWCP (B4) | yes | 0.810 | 0.809 | 0.798 | 0.803 | 0.989 | 0.937 | 0.253 | 0.000 |
| ProtCOVER | no | 0.807 | 0.800 | 0.765 | 0.820 | 0.978 | 0.936 | 0.261 | 0.001 |
| ProtCOVER | yes | 0.822 | 0.822 | 0.815 | 0.812 | 1.020 | 0.925 | 0.234 | 0.001 |
| ProtCOVER-D | no | 0.807 | 0.793 | 0.760 | 0.824 | 0.957 | 0.935 | 0.264 | 0.000 |
| ProtCOVER-D | yes | 0.808 | 0.810 | 0.798 | 0.812 | 0.984 | 0.946 | 0.263 | 0.000 |
| ProtCOVER-W | no | 0.808 | 0.803 | 0.766 | 0.820 | 0.979 | 0.948 | 0.251 | 0.000 |
| ProtCOVER-W | yes | 0.815 | 0.811 | 0.796 | 0.813 | 1.000 | 0.937 | 0.248 | 0.000 |
| (i)+(iii) gate | no | 0.797 | 0.791 | 0.753 | 0.810 | 0.958 | 0.938 | 0.279 | 0.002 |
| (i)+(iii) gate | yes | 0.812 | 0.813 | 0.804 | 0.800 | 1.001 | 0.930 | 0.249 | 0.004 |
| ProtCOVER-W + gate | no | 0.808 | 0.803 | 0.767 | 0.821 | 0.982 | 0.945 | 0.250 | 0.002 |
| ProtCOVER-W + gate | yes | 0.816 | 0.812 | 0.797 | 0.814 | 1.008 | 0.930 | 0.246 | 0.005 |
| Support-only (B5) | no | 0.808 | 0.803 | 0.767 | 0.821 | 0.981 | 0.947 | 0.250 | 0.002 |
| Support-only (B5) | yes | 0.816 | 0.812 | 0.797 | 0.814 | 1.005 | 0.933 | 0.245 | 0.005 |
| Random abst. (B5) | no | 0.808 | 0.803 | 0.767 | 0.821 | 0.981 | 0.946 | 0.250 | 0.002 |
| Random abst. (B5) | yes | 0.816 | 0.812 | 0.797 | 0.814 | 1.006 | 0.933 | 0.246 | 0.005 |
